# GNPS Dashboard: Collaborative Analysis of Mass Spectrometry Data in the Web Browser

**DOI:** 10.1101/2021.04.05.438475

**Authors:** Daniel Petras, Vanessa V. Phelan, Deepa Acharya, Andrew E. Allen, Allegra T. Aron, Nuno Bandeira, Benjamin P. Bowen, Deirdre Belle-Oudry, Simon Boecker, Dale A. Cummings, Jessica M Deutsch, Eoin Fahy, Neha Garg, Rachel Gregor, Jo Handelsman, Mirtha Navarro-Hoyos, Alan K. Jarmusch, Scott A. Jarmusch, Katherine Louie, Katherine N. Maloney, Michael T. Marty, Michael M. Meijler, Itzhak Mizrahi, Rachel L Neve, Trent R. Northen, Carlos Molina-Santiago, Morgan Panitchpakdi, Benjamin Pullman, Aaron W. Puri, Robin Schmid, Shankar Subramaniam, Monica Thukral, Felipe Vasquez-Castro, Pieter C Dorrestein, Mingxun Wang

## Abstract

Access to web-based platforms has enabled scientists to perform research remotely. A critical aspect of mass spectrometry data analysis is the inspection, analysis, and visualization of the raw data to validate data quality and confirm statistical observations. We developed the GNPS Dashboard, a web-based data visualization tool, to facilitate synchronous collaborative inspection, visualization, and analysis of private and public mass spectrometry data remotely.

## Maintext

Web-based computing has changed our digital lives. The recent disruptions to office and laboratory workspaces resulting from the COVID-19 pandemic, including campus closures, telework, and stay-at-home orders, all increased the need for the development of novel online approaches to scientific research. In particular, the responses to these needs revealed that near-real-time synchronous interactive web applications used simultaneously by more than one person in the same analysis environment (similar to collaborative text editing in Google Docs or Microsoft Office 365), would enable a level of collaborative research that was simply not possible in traditional work environments.

One of the key analytical chemistry techniques applied to the life sciences is mass spectrometry (MS). Although there are numerous software solutions for analysis of MS data, most require the installation of specific software packages, advanced knowledge in command-line based execution, expert knowledge of where data can be found, downloading of data with file transfer protocols to a local drive, and conversion of data into compatible formats^1–3^. Notably, many MS data analysis software packages suffer from incompatibility with different data formats and poor interoperability. The limited accessibility of analysis software poses a significant barrier for MS data inspection by both experts and non-experts, which prevents collaborators, reviewers, and readers of scientific publications from inspecting the raw MS data to reproduce and verify interpretation or discuss data while inspecting it. Although current web-based visualizations of MS data are available for single spectrum visualization^4–7^, there are no open browser visualization solutions for full MS datasets. With the rapid growth in MS data availability^6,8–10^ and the potential to leverage large MS datasets to develop novel hypotheses, it is becoming increasingly important for the scientific community to have open and transparent solutions to share, inspect, and reproduce MS data and its analysis.

We developed the GNPS Dashboard, a centralized web resource (https://gnps-lcms.ucsd.edu), to facilitate the visualization of liquid and gas chromatography-mass spectrometry (LC-MS and GC-MS) data for quality inspection, visualization, sharing, collaborative examination, and hands-on teaching of MS concepts using private and publicly available MS data, including files stored in the MS data repositories GNPS/MassIVE^6^, MetaboLights^9^, ProteomeXchange^11^, and Metabolomics Workbench^8^ (**Fig. 1**, [Link to Instructions](https://ccms-ucsd.github.io/GNPSDocumentation/lcms-dashboard/)). All publicly shared MS files from compatible repositories can be viewed, selected, and compared in the GNPS dataset explorer (https://gnps-explorer.ucsd.edu/). Files not deposited in public MS repositories can be visualized through a drag-and-drop option for file transfer. Although .mzXML, .mzML^12^, .CDF, and Thermo .raw file formats are compatible with GNPS Dashboard and can be directly uploaded for analysis **(SI Use Case 9**), GNPS’s quickstart interface (https://gnps-quickstart.ucsd.edu/conversion) or Proteowizard^13^ should be used to convert files to a compatible format. Via deep linking from the GNPS platform, GNPS Dashboard serves as a data explorer and central hub for further data analysis from Classical Molecular Networking^6^ (**SI Use Case 9**) and Feature-based Molecular Networking^14^ (**SI Use Cases 1 and 8**), MSHub GC-MS deconvolution^15^, *in silico* spectrum annotation via SIRIUS and CSI:FingerID^16^ (**SI Use Case 10**), and MASST^17^ (**SI Use Case 5**).

**Figure 1:**
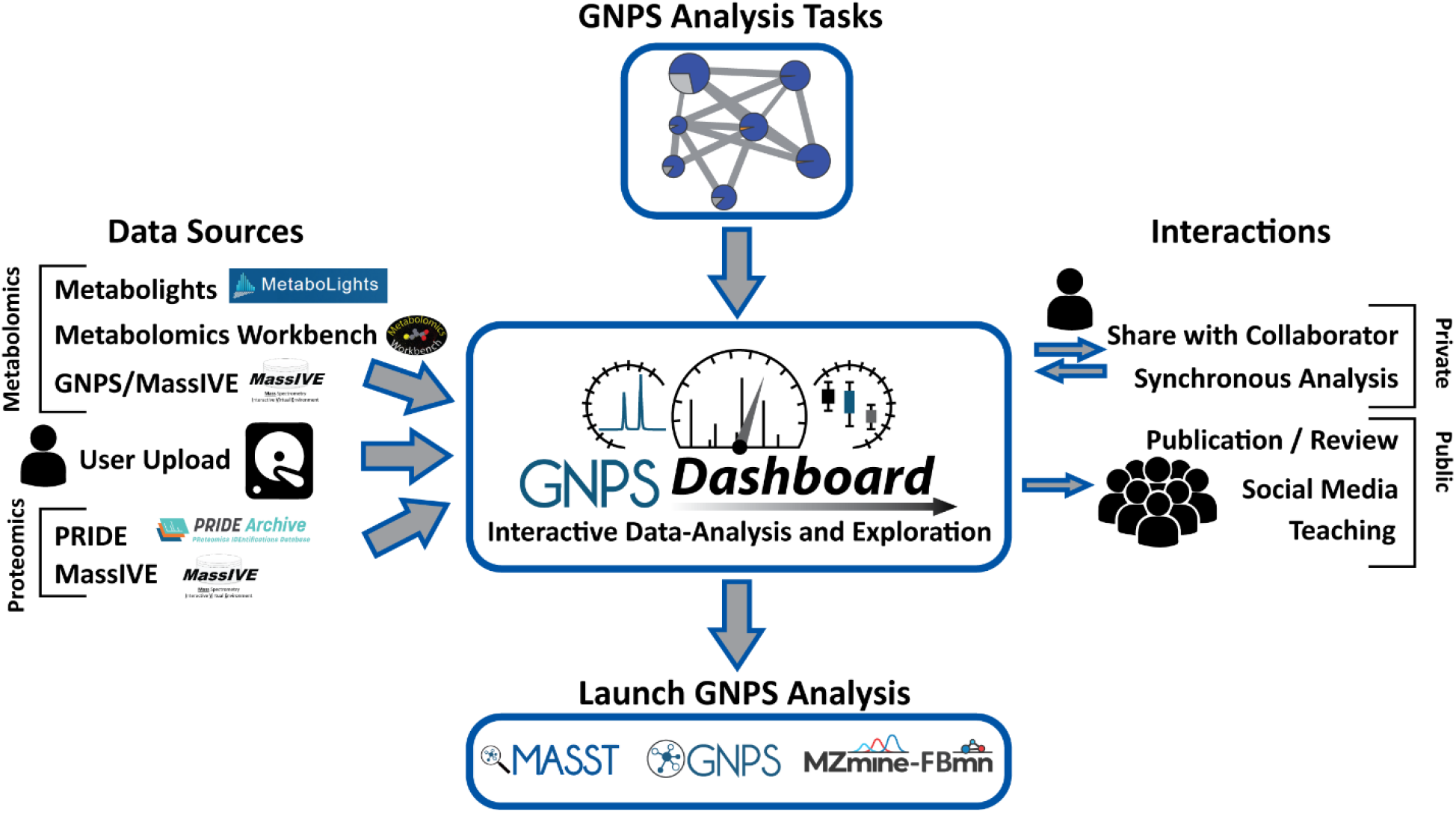
Overview of the GNPS Dashboard in the Data/Analysis Ecosystem. The GNPS Dashboard has been integrated into the web-accessible mass spectrometry ecosystem. The top and left panels show how the GNPS Dashboard can ingress datasets from public metabolomics resources: MetaboLights, Metabolomics Workbench, and GNPS/MassIVE, proteomics resources: PRIDE and MassIVE, private user uploads, and private GNPS analysis tasks. The bottom panel shows that directly out of the GNPS Dashboard, users can launch downstream analysis on their data. Finally, the right panel shows how the GNPS Dashboard and its visualizations can interact with the wider community with reproducible URL or QR code link outs from any analysis and teaching/synchronous collaboration modes for real-time interactivity.

The GNPS Dashboard enables rapid inspection of Total Ion Chromatograms (TIC), 2D retention time versus *m/z* heat map for global inspection of all signals, Extracted Ion Chromatograms (XIC) (**SI Use Cases 1, 2, 12, and 15**), and tandem mass spectra (MS/MS) (**SI Use Case 13**) for inspection/visualization of individual compounds, as well as quantitative comparison of the peak abundances of two groups as box-plots (**Fig. 2 and SI Use Case 7)**. Publication-quality figures of each display item are automatically generated for download in scalable vector graphic (.svg) format. Further, the GNPS Dashboard can aid peer review of scientific manuscripts (**SI Use Case 17**) and inspecting public quantitative proteomics data to validate published results (**SI Use Cases 6 and 13**). Beyond visualization and analysis of MS data, the GNPS Dashboard has been shown to support the development of other bioinformatics tools (**SI Use Case 14**) that may not have their own web-enabled user interfaces.

**Figure 2:**
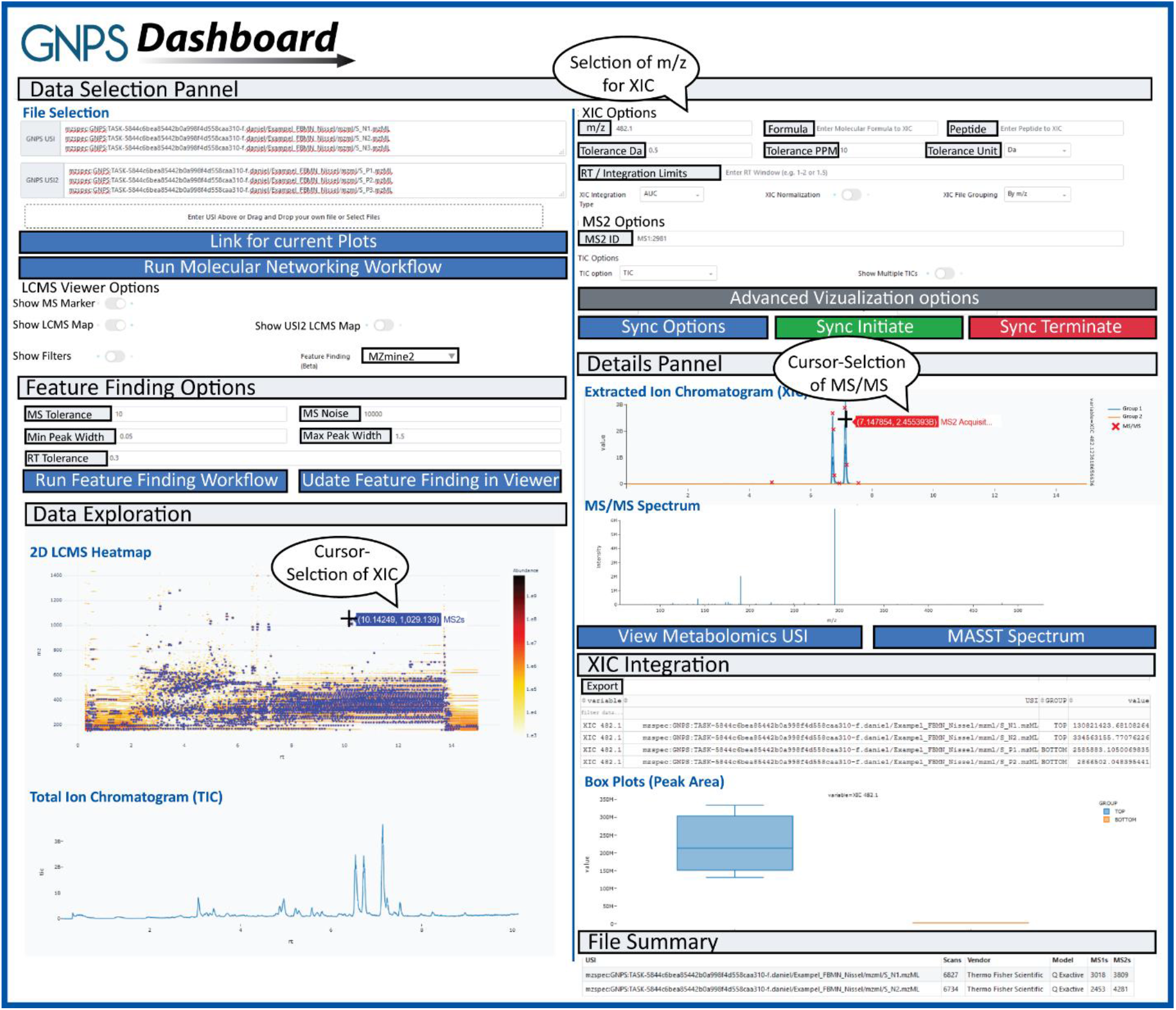
Overview of the GNPS Dashboard User Interface: The GNPS Dashboard’s user interface includes data selection and data exploration visualization via 2D LC-MS heatmaps, total ion chromatogram, extracted ion chromatogram (XIC) and tandem mass spectra as well as simple feature finding options and manual XIC integration. Any analysis and visualization that is created within the GNPS Dashboard can be shared as a URL or quick response (QR) code for the community to visualize, reproduce, and improve the data analysis.

The GNPS Dashboard encodes the full state of the visualization interface in an easily-shareable link, which empowers users to share the exact same visualization with others (e.g., including pan/zoom configurations), thus reducing miscommunication and improving data transparency, for example during (remote) meetings with collaborators (**SI Use Case 3**). Every visualization and analysis result can be shared via a URL that will re-launch the original data visualization on their device along with the history of the analysis (enabling users to rewind and step forward each discrete analysis step, up to 1000 steps per session). Users can share these links with collaborators and embed them in publications, presentations, or social media posts (e.g. [Example Tweet](https://twitter.com/mingxunwang/status/1356757390863912961) and **SI Use Case 4**). As with the shared link, a final visualization can also be shared as a Quick Response (QR) code. Anyone with a link or QR code can build upon the analysis and re-share their additions (**SI Use Case 3**). Links and QR codes will remain valid, accessible, and embeddable in publications and presentations for data that has been archived in a public repository.

To further and uniquely enhance remote collaborations or classroom teaching, the GNPS Dashboard includes leader-follower synchronization (real-time updates from one user) and fully collaborative synchronization (real-time updates from multiple users). The leader-follower mode enables followers to mirror a leader’s analysis in real-time. The followers can then disconnect the synchronization at any time to continue the analysis from where the leader left off without needing to reload the data. Synchronization modes facilitate collaborations and remote as well as in-person teaching. The GNPS Dashboard leader-follower paradigm has already been used in remote classroom teaching by at least five institutions, including undergraduate institutions, with up to 50 students per classroom (**SI Use Case 5**). The fully collaborative synchronization enables multi-user simultaneous shaping of the visualization and data exploration in a manner similar to online synchronous collaborative document editing (**SI Use Case 18**). For example, users can initiate a collaborative session with two or more people on any web-accessible device and can edit simultaneously with this link (GNPS Collab Start [Link](https://gnps-lcms.ucsd.edu/?synchronization_type=COLLAB) and [Instructions](https://ccms-ucsd.github.io/GNPSDocumentation/lcms-dashboard/#fully-sychronized-visualization-session)). In these synchronization and standard analysis modes, not only is the final state of the analysis saved but so is every discrete action, enabling users to follow advancement or review past evolution of data analysis. A snapshot and history of the collaborative work can be created and shared.

While the GNPS Dashboard is accessible as a free public web service, it is possible to locally install the GNPS Dashboard to function with local data sources, making collaborative analysis and sharing possible, privately, within an institution when necessary (*e*.*g*. government agencies and clinical laboratories). Although only recently introduced, the GNPS Dashboard has already supported the visualization of 8,144 mass spectrometry files from a worldwide user base (October 2020 - April 2021). We envision that over time new features will be added to the GNPS Dashboard to support new mass spectrometry data types (*e*.*g*., ion mobility) and visualizations, in collaboration with the mass spectrometry community.

Overall, the GNPS Dashboard facilitates the visualization and exploration of mass spectrometry data online, which significantly lowers the barriers to entry for data analysis, and introduces new modes of collaborative data analysis. Tying the capabilities of the GNPS Dashboard together with public data repositories will improve public data analysis, promote re-analysis, encourage data transparency and sharing, and strengthen the reproducibility of data analysis.

## Online Methods

The GNPS Dashboard itself is the intersection of several web and mass spectrometry technologies and standards, including the Universal Spectrum Identifier (USI)^7^, pymzML^18^, ProteoWizard^13^, ThermoRawFileParser^19^, Dinosaur^20^, and MZmine^2^. The dashboard is built on web technologies with a Python backend and the inclusion of mass spectrometry-specific tools to enable automatic conversion of data and access to the data. The open source data science tools DataShader (Holoviz) and Plotly/Dash (Plotly Inc.) are used for visualization. Finally, the dashboard is integrated into public mass spectrometry resources with the ability to resolve, translate, and acquire data via the mass spectrometry USI.

The run environment and deployments are handled via Docker and Docker-Compose. A full set of dependencies and configurations are written into the accompanying source files for the GNPS Dashboard. Detailed documentation on how to use the GNPS dashboard can be found here ([Link](https://ccms-ucsd.github.io/GNPSDocumentation/lcms-dashboard/)).

## Data availability

All data used within this manuscript and supplemental use cases is publicly available through the MassIVE Repository (massive.ucsd.edu) and proteomeXchange (proteomexchange.org) under the following accession numbers: MSV000086206, MSV000086834, MSV000082378, MSV000086996, MSV000086079, MSV000082493, MSV000085618, MSV000085852, MSV000086584, MSV000086453, MSV000085974, MSV000085376, MSV000086729, MSV000081885, MSV000085070, MSV000086092, MSV000079843/PXD015300, MSV000083508/PXD010154, MSV000087056, MSV000087075, MSV000083859, MSV000087157.

The processed data highlighted in each use case can be downloaded and visualized in the included urls in each example in the supplemental information.

## Code availability

The GNPS Dashboard and GNPS Dataset Explorer and their source code are available through the GNPS web environment and GitHub, enabling quick installation on local servers.

The GNPS Dashboard source code can be found on GitHub: https://github.com/mwang87/GNPS_LCMSDashboard under a modified UCSD BSD License.

The GNPS Dataset Explorer source code can be found on GitHub: https://github.com/mwang87/GNPS_DatasetExplorer under an MIT License.

## Acknowledgments

This work was, in part, supported by the National Institutes of Health (NIH) with grant numbers U19AG063744, U2CDK119886, OT2 OD030544, GM107550, R03CA211211, R24GM127667, 1R01LM013115 and P41GM103484, the National Science Foundation (NSF) with grant IOS-1656475 and ABI 1759980, and the Gordon and Betty Moore Foundation (GBMF7622). VVP was supported by the L.S. Skaggs Professorship and Therapeutic Innovation Award from the ALSAM Foundation and NIH R35GM128690. NB and BP were supported by NIH P41 GM103484. MTM was supported by NSF grant CHE-1845230. DB was supported by NSF grant DUE 16-25354.The I.M laboratory was supported by grants from the European Research Council (No. 640384) and from the Israel Science Foundation (ISF No. 1947/19). TRN, BB, and KL were supported by the U.S. Department of Energy Joint Genome Institute, a DOE Office of Science User Facility, which is supported by the Office of Science of the U.S. Department of Energy under Contract No. DE-AC02-05CH11231.

Furthermore, we would like to thank Tristan de Rond, Laura-Isobel McCall, Wout Bittremieux, Kelly Weldon, and Emily Gentry for testing the software and suggesting updates. We would like to thank Vagisha Sharma, for working on the standalone MSView app during her masters degree research 12 years ago, which provided some visualization inspiration for the GNPS Dashboard. We thank Claire O’Donovan and the team at MetaboLights for developing a well-documented API. AWP and DACJ thank N. Cecilia Martinez-Gomez (UC Berkeley) for *Methylorubrum extorquens* PA1 Δ*cel* strain CM2730, and Ming Hammond (University of Utah) for use of her LC-MS system. Lastly, we thank all members of the research community who make their data publicly accessible, which contributes to open, transparent, and reproducible science.

## Author Contributions

MW and DP conceived the project. MW developed the software for GNPS Dashboard and GNPS Dataset Explorer. MW, DP, VVP, and PCD provided guidance and supervision. MW, DP, VVP, KNM, ATA, AKJ, BP, DACJ, AWP, CMS, MT, NG, NB, and PCD tested and offered user feedback for the GNPS Dashboard. SS and EF implemented the APIs in Metabolomics Workbench to support GNPS Dashboard. TN, BB, KL facilitated making metabolomics data in Genome Portal available in GNPS Dashboard. MW, DP, VVP, KNM, ATA, AKJ, SAJ, DA, BP, DACJ, AWP, CMS, MT, AEA, RS, NB, SB, MTM, DBO, MNH, RG, MMM, IM, JH, FVC, JMD, NG, NB, RLN, and PCD wrote use cases for the GNPS Dashboard Supplemental Information. PCD provided funding for the project. PCD, MW, VVP, and DP wrote the draft manuscript. All authors edited and approved the final manuscript.

