## Supplemental Data 1 for "GNPS Dashboard: Collaborative Analysis of Mass Spectrometry Data in the Web Browser"

### Protocol: Exploring LCMS-MS data

#### Part I: Determining Exact Mass using ChemDraw

1. Navigate to the [PLNU virtual desktop (https://view.pointloma.edu/)](https://view.pointloma.edu/).
    - Select the **VMware Horizon HTML Access** option.
    - Login with your PLNU credentials.
    - Select "Computer Lab Student."
2. Save the .cdx file onto the virtual desktop.
    - ***From within the virtual desktop,*** open a browser, and log into Canvas. From in Canvas on the virtual desktop, navigate to this page and open this link: [$WIKI_REFERENCE$/pages/chemdraw-files-for-week-9-lcms (https://canvas.pointloma.edu/courses/56791/pages/chemdraw-files-for-week-9-lcms)](https://canvas.pointloma.edu/courses/56791/pages/chemdraw-files-for-week-9-lcms) (https://canvas.pointloma.edu/courses/56791/pages/chemdraw-files-for-week-9-lcms)
    - 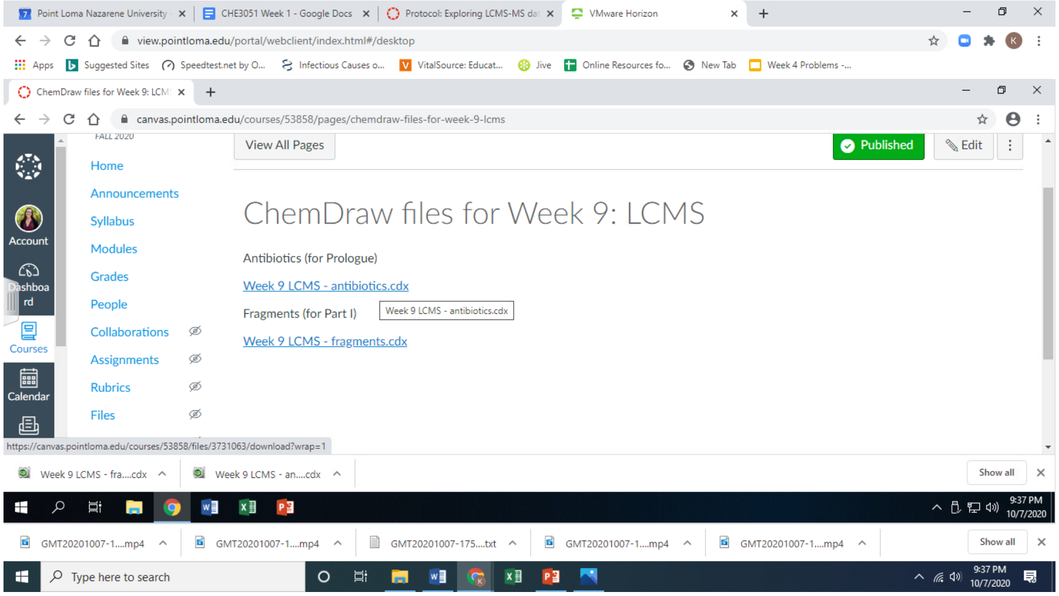

    - Download both .cdx files and note where they save (probably the Downloads folder).
    - *Note that you have to be* ***in the virtual desktop window*** *when downloading. If you try to do it from another tab in the browser on your computer, you'll be saving the file to your computer, instead of to the computer lab computer.*
3. Open the file in ChemDraw.
    - From *within the virtual desktop,* Click on the Start Menu icon to open the start menu, and select ChemOffice 2019, followed by ChemDraw 19.1.

- 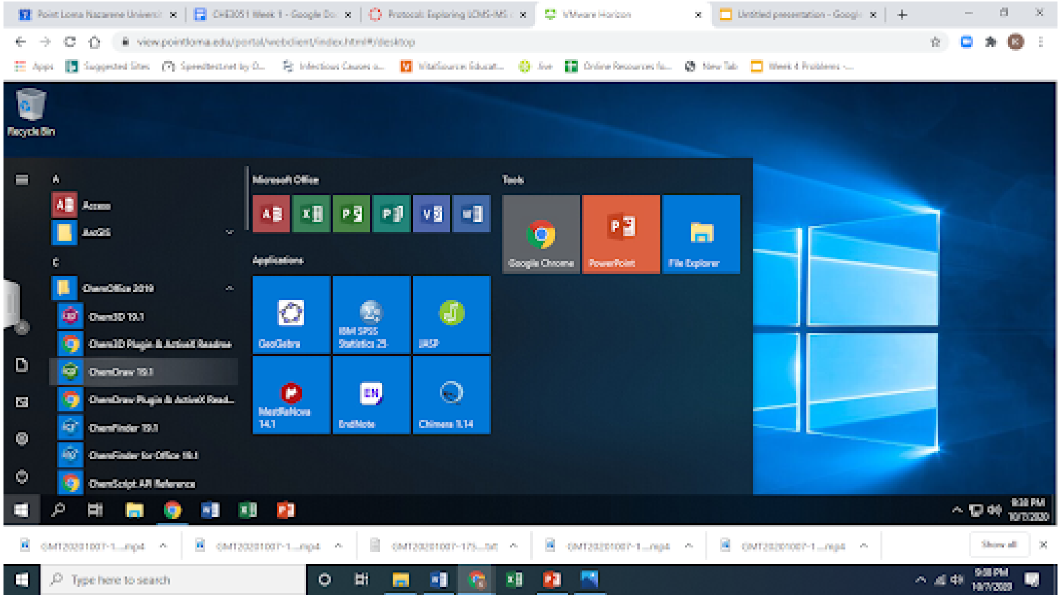
- In the menu at the top of the ChemDraw screen, click File, then Open, then navigate to the file.

4. 'Ionize' the antibiotic structures. Recall that the mass spectrometer doesn't detect neutral molecules, but charged ions - in this case $(M\text{-}H)^-$ ions, also known as the conjugate base.
    - From the toolbar on the left side, click on the Chemical Symbols tool (the default is usually ⊕) and hold the button down to see the full set of options.
    - 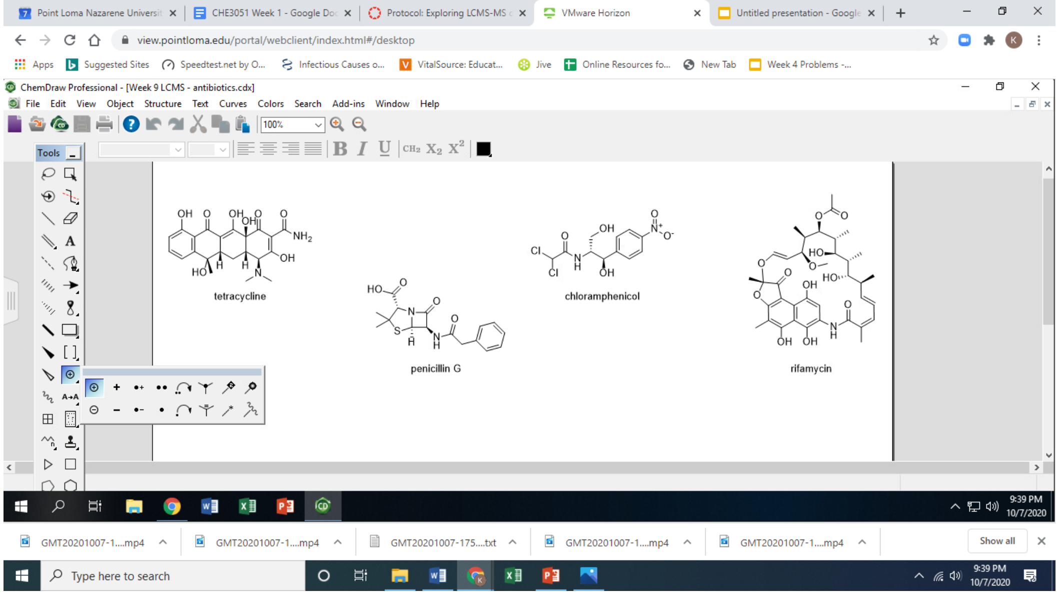

    - Select the negative charge (⊖).
    - Hover your cursor over the oxygen on one of the OHs on tetracycline, until the O is highlighted. Click once to impart a negative charge on this oxygen. (The hydrogen will disappear.)
    - Repeat for one of the OH groups in each of the other antibiotics.
5. Check the masses you determined in your pre-lab using ChemDraw.
    - In the top menu, click View, then make sure Show Analysis Window is checked.
    - Using the lasso tool (top left in the toolbar), select one of the antibiotics.
    - The molecular formula, exact mass, molecular weight, etc. will appear in the Analysis Window.

- 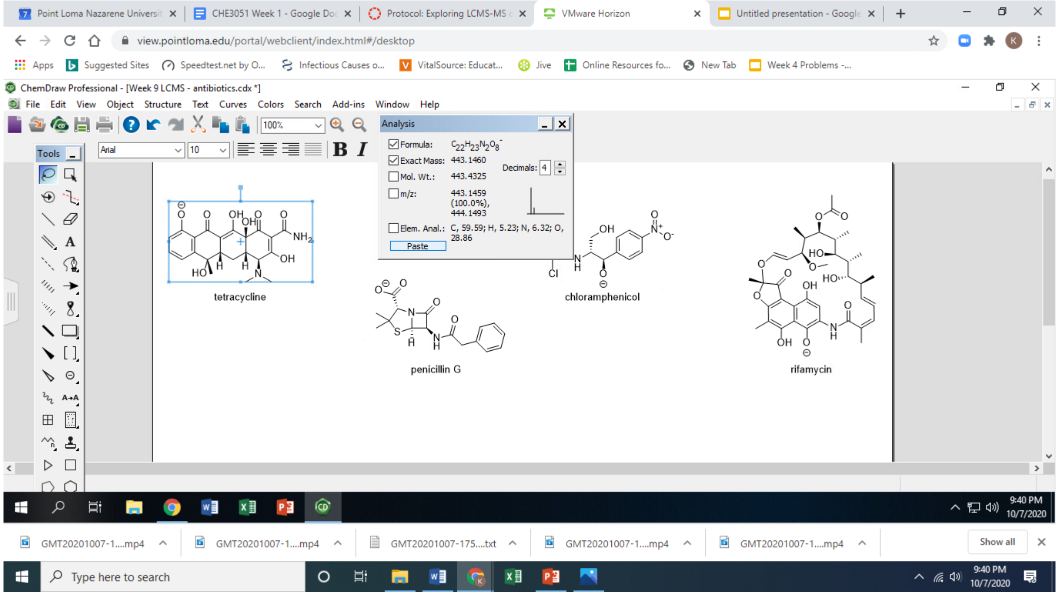

- Adjust the number of Decimals to 4. Uncheck the boxes next to Mol. Wt., m/z:, and Elem. Anal. Then click Paste.
- Repeat with the other antibiotics. How do these values compare with the ones you calculated?
- You can minimize the virtual desktop for now, but don't close it. (You'll need to use ChemDraw again in step 8 of Part II.)

#### Part II: Exploring the LCMS of a mixture of antibiotics

1. Visit gnps.ucsd.edu (https://gnps.ucsd.edu/ProteoSAFe/static/gnps-splash.jsp) and log into your GNPS account
   - 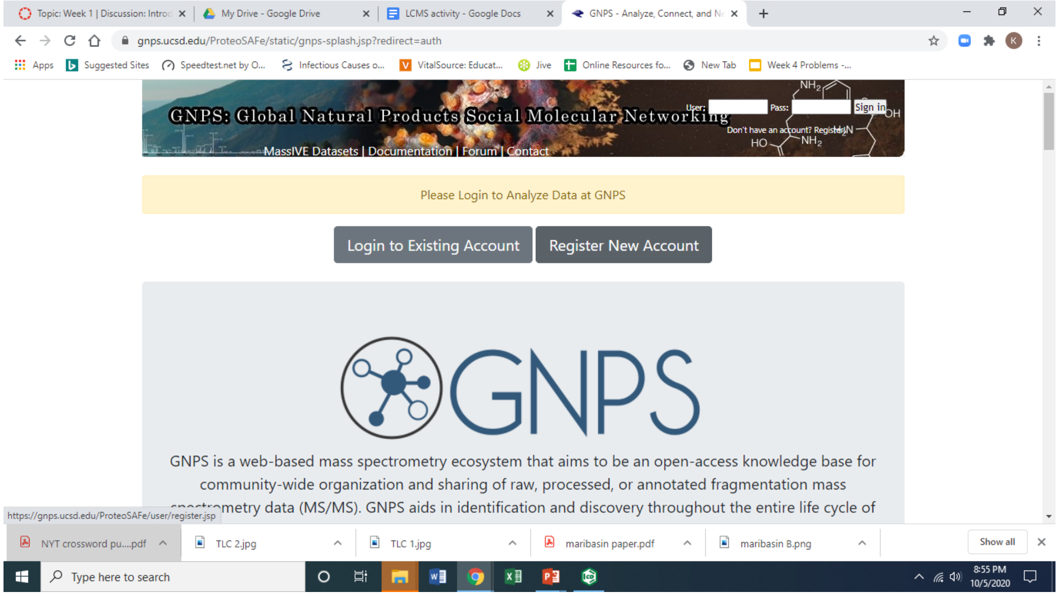

2. In a separate window, go to the GNPS LCMS Dashboard. (https://gnps-lcms.ucsd.edu/) LCMS Browser website (https://gnps-lcms.ucsd.edu/?usi=mzspec%3AMSV000086079%3Aantibiotics_1ug_ml_neg_20200902112656%3Ascan%3A1000&usi2=&xicmz=&xic_tolerance=0.5&xic_rt_window=&xic_norm=False&xic_file_grouping=FILE&show_ms2_markers=True&ms2_identifier=None)

3. Open the antibiotic mixture datafile. Under the **File Selection** tab, in the **GNPS USI** field, delete the default file path and paste the following: **mzspec:MSV000086079:antibiotics_1ug_ml_neg_20200902112656:scan:1000**, then click **Link to these plots**.
    - 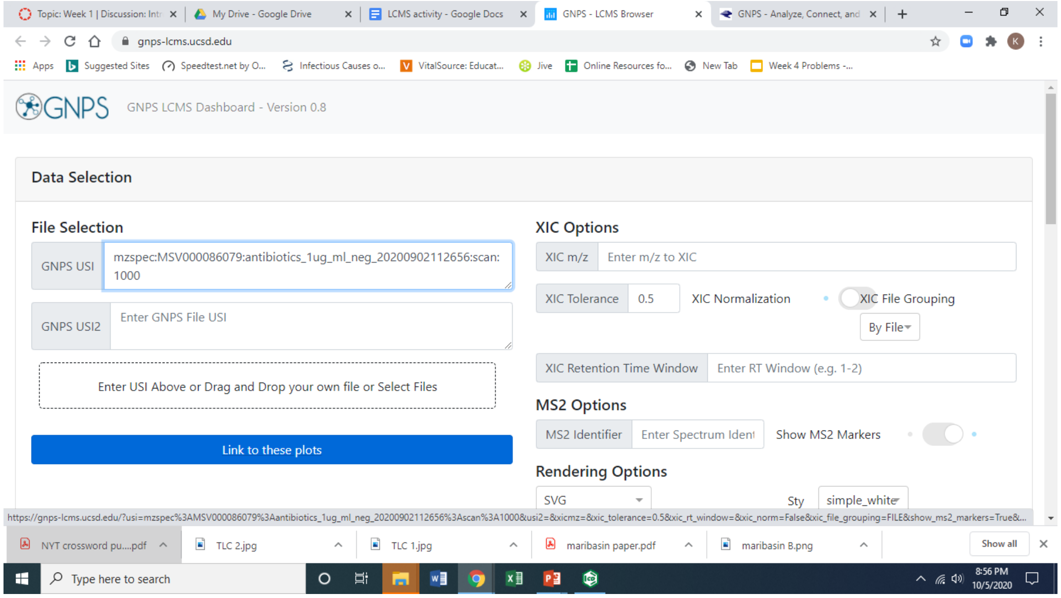

4. Save an image of the TIC plot. Scroll to the bottom of the page (left side). If you hover your cursor over the TIC plot, a set of icons will appear directly above, including one that looks like a camera. Click on this icon to download an image (as an SVG file) of your TIC to include with your postlab report.
    - 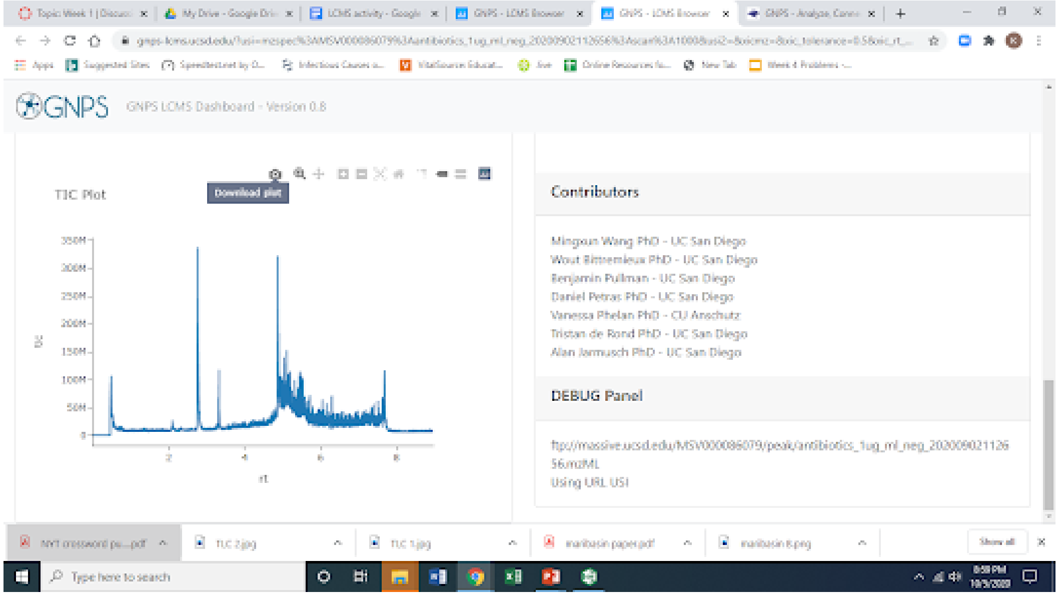

5. Now let's explore the **TIC plot.** The **TIC** - or **Total Ion Chromatogram** - is a plot of the total number of ions (of all masses) hitting the detector at a given time. The y-axis shows the number of ions, while the x-axis shows the **retention time (rt)** in minutes. When a molecule such as an antibiotic is coming off the chromatography column, the total number of ions goes up, and then down as the compound tapers off. In this plot, each peak corresponds to a molecule.
    - Hover your cursor over a peak, such as the tallest one in the spectrum, between 2 and 4 minutes. When you are near a peak, a little box will pop up with the exact retention time (rt) and total ion counts (tic) for that peak.

- 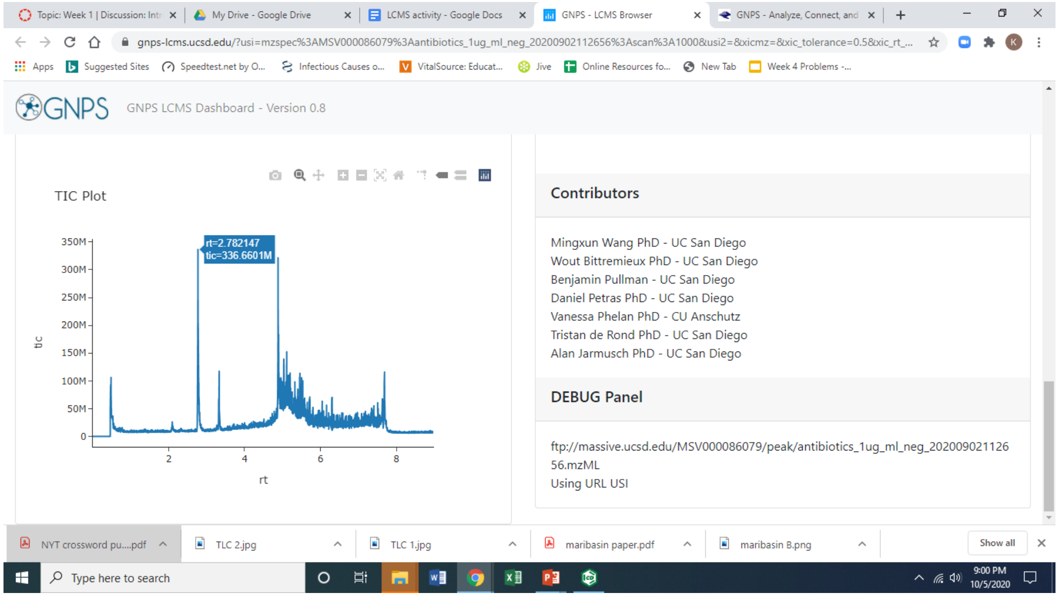

- Click on the peak at ~2.8 min. Wait a few seconds, while the mass spectrum (MS1) loads to the right of the TIC plot. The **MS1** (also just known as the **mass spectrum**) shows each of the masses detected at a given time (in this case 2.8 minutes). The x-axis shows the mass of each ion (m/z), while the y-axis shows the intensity (number of ions hitting the detector) corresponding to each mass. A typical mass spectrometry dataset consists of *hundreds* or *thousands* of mass spectra like this one - one corresponding to each datapoint in the TIC plot!
- Now scroll right and hover your cursor near the top of the large peak in the MS1. Another little box will appear, showing the mass of that peak. This is the molecular ion for the antibiotic that came off at 2.8 minutes. Based on the calculations you did for your pre-lab, which antibiotic is this?
- 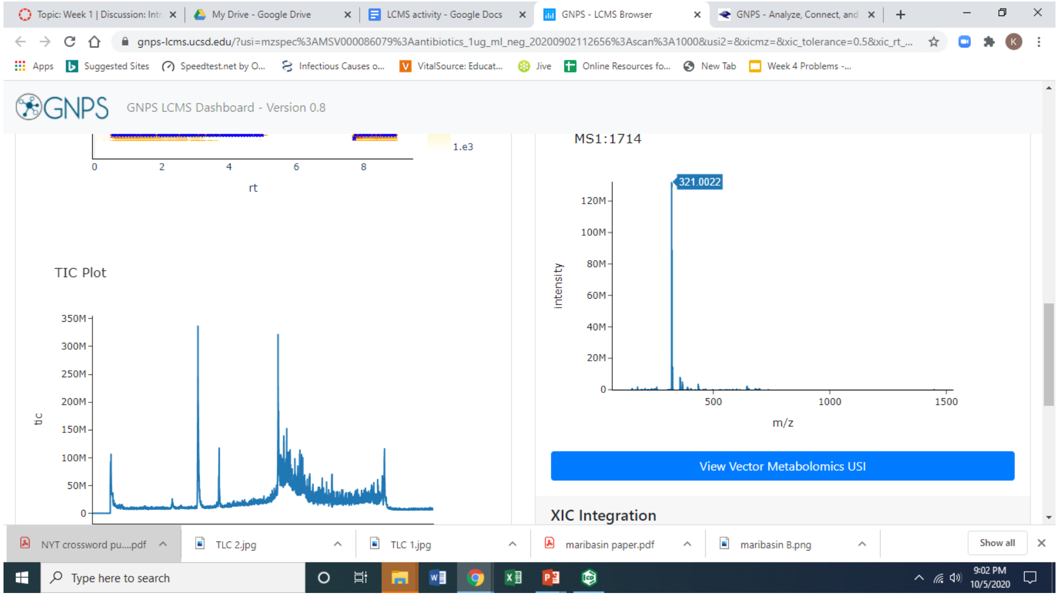


6. Find each antibiotic in the TIC plot. Repeat step 7 for the peaks at 2.1 (small peak), 3.3 (medium-sized peak) and 4.9 minutes. Which antibiotic does each correspond to? Download an image of one of the MS1 spectra to include with your postlab report.
7. Let's explore something called the **XIC**. In this example, our total ion chromatogram (TIC) is quite 'clean'. There were only a few purified antibiotics in this sample, and these show up as distinct peaks in the chromatogram. In other cases - such as when we are looking at natural product

extracts - the TIC may be packed with peaks, most of which we are not interested in. In that case, we can quickly pull out peaks of interest using something called an **extracted ion chromatogram** (or **XIC**).

- Scroll back to the top of the page. Under **XIC Options**, type the exact mass of the conjugate base of penicillin (the one given to you in your prelab): 333.0915. Scroll down and wait a few seconds for the XIC Plot to appear under the Details Panel. The XIC Plot will show *only* the ions that have an exact mass of 333.0915. What does the XIC Plot look like?
- 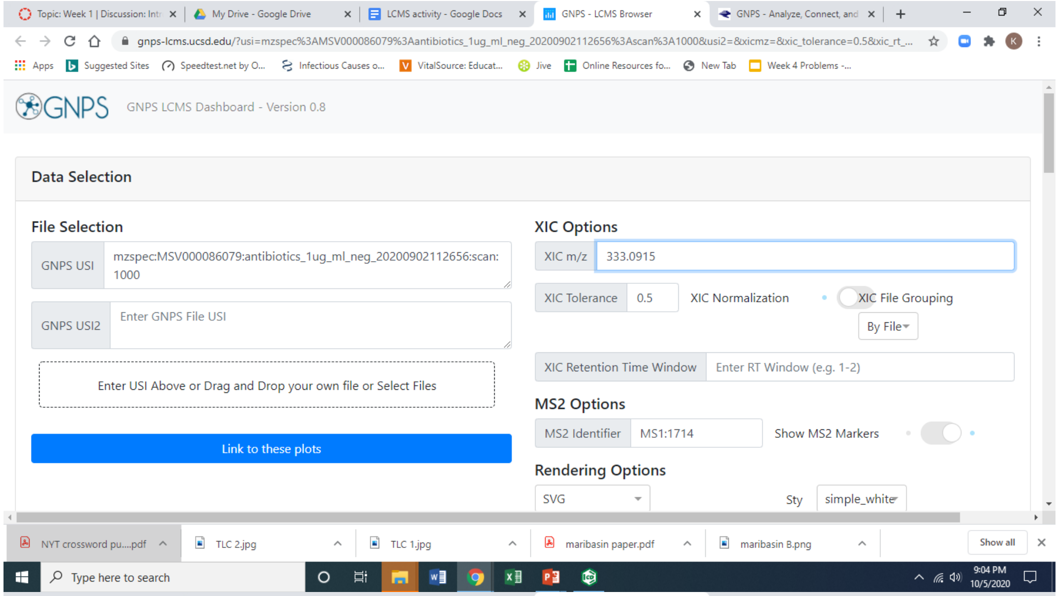

- 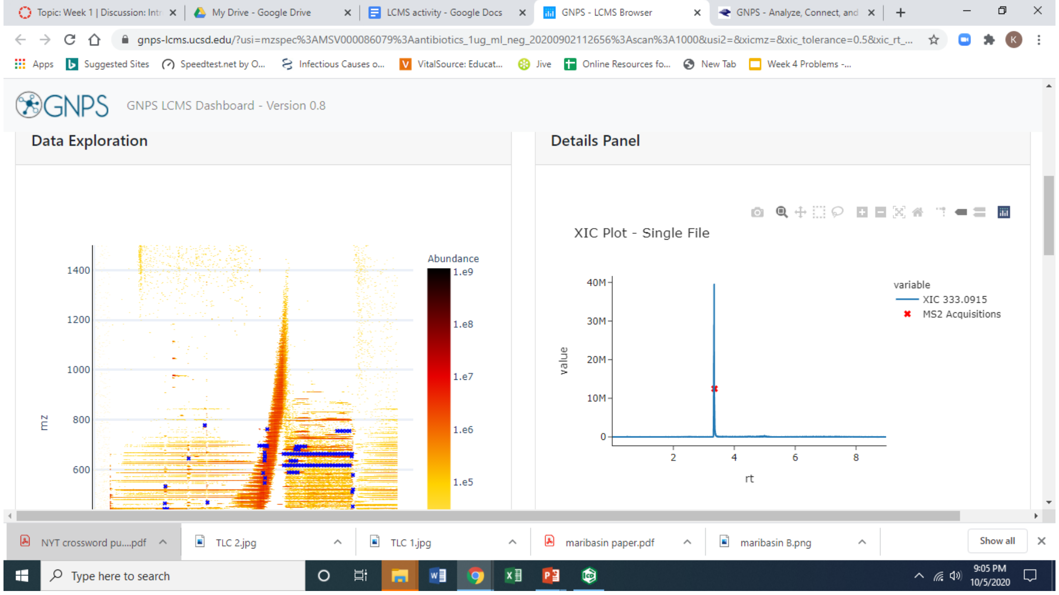

- In the XIC Options field, add a semicolon after the mass of penicillin, and add the other masses you calculated in your prelab for each of the antibiotics, separated by semicolons. How does this plot compare to the TIC plot?
- Download an image of the XIC plot of the four antibiotics to include with your postlab report.

8. Now let's explore the tandem mass spectrum (MS2 spectrum).
    - At the top right of the page, next to **MS2 Options**, check that the **Show MS2 Markers** toggle switch is toggled to the right ('on'). If not, press on the toggle switch once. (When Show MS2

Markers is ‘on’, you should see a series of tiny blue Xs in the 2D plot under **Data Exploration**.)

- 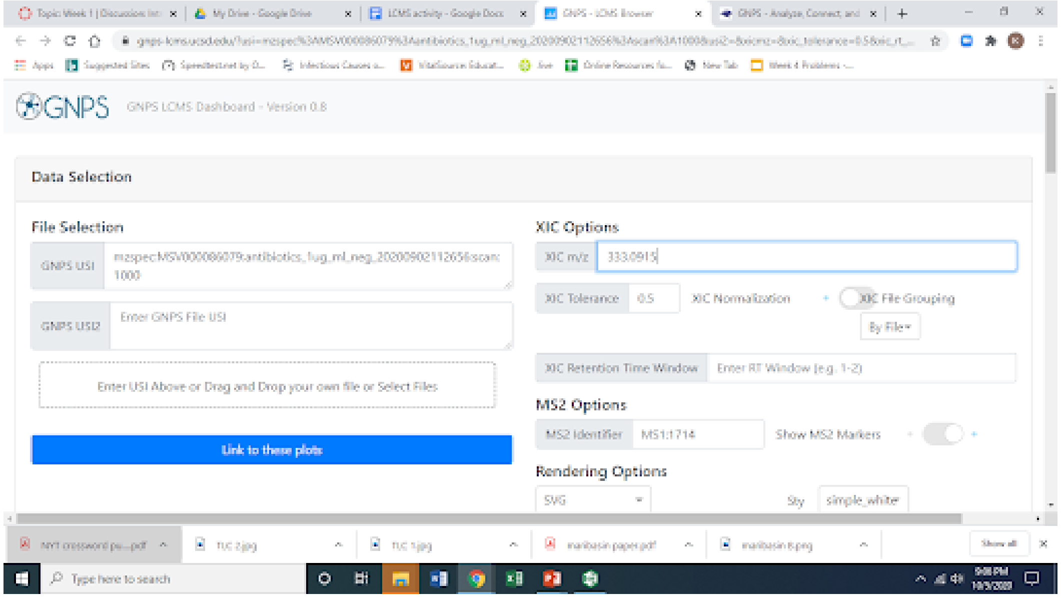
- Scroll down to the 2D plot under the Data Exploration heading. This somewhat overwhelming 2D plot shows the combined information from all the MS1 spectra in this dataset! The x-axis shows retention time, while the y-axis shows mass-to-charge ratio. Each vertical slice in the plot corresponds to the MS1 spectrum at that retention time. If you look at the retention times corresponding to our antibiotics, you can see vertical stripes that look different from the general streakiness of the plot. Each blue X in the plot corresponds to a particular ion (peak) in a particular MS1 spectrum (vertical slice) that was selected for MS2.
- 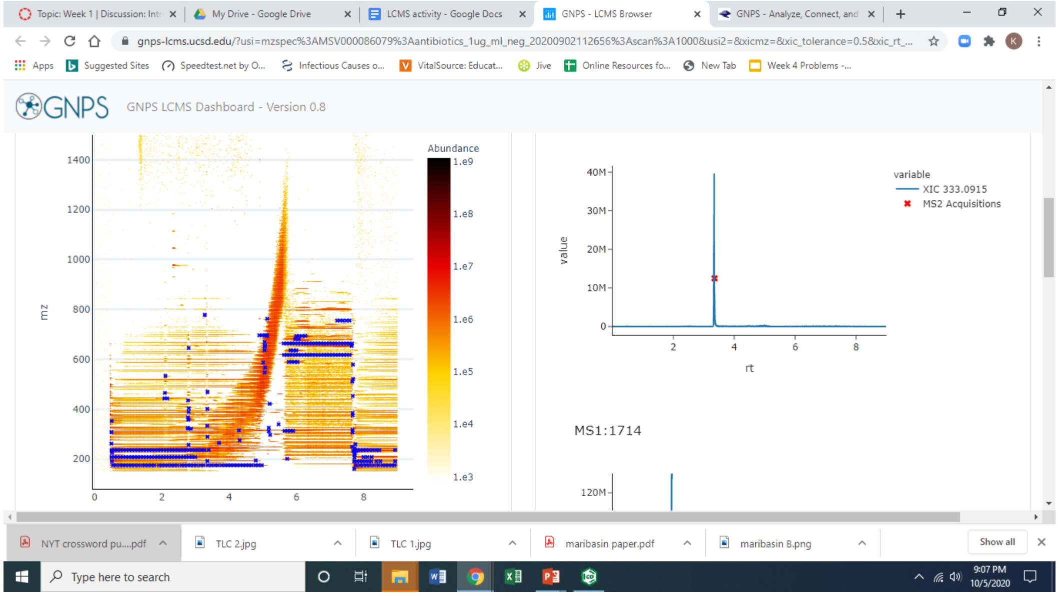
- Display the MS2 spectrum for tetracycline. In the Data Exploration 2D plot, move your cursor around to find the blue X at the retention time and mass for tetracycline that you identified in step 8. When you find the corresponding blue X (at rt ~2.16 min, m/z 443.1428), click on it and wait a few seconds for the Details Panel to update. Scroll down to see the MS2 spectrum that opens.

- 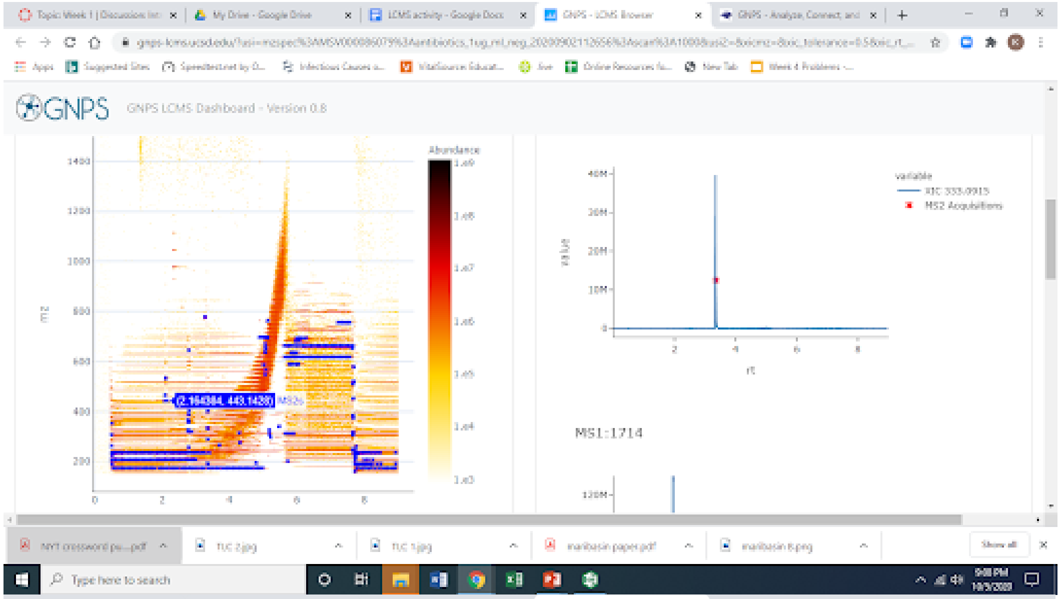
- 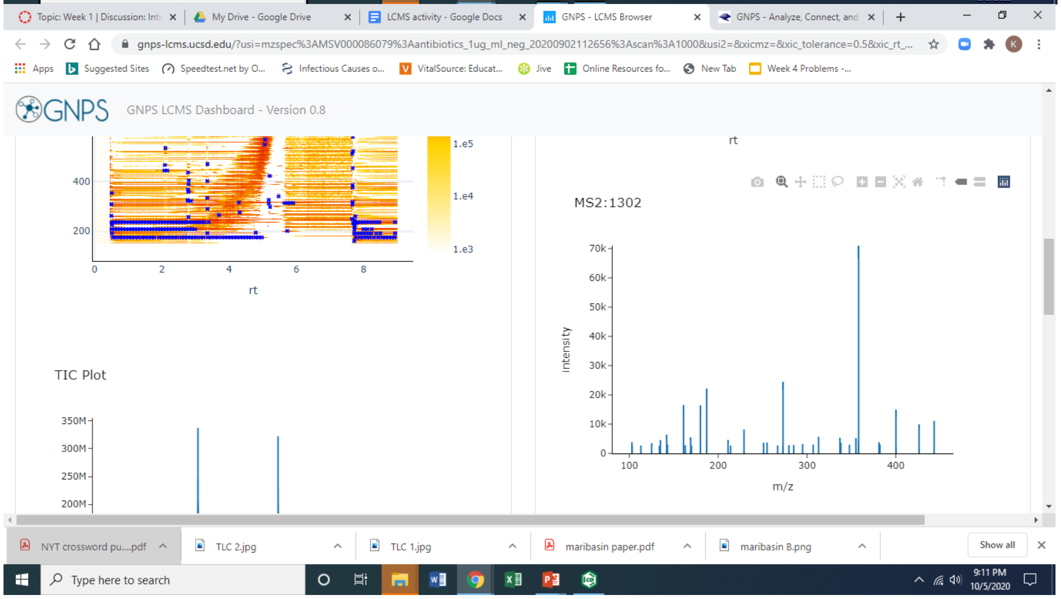
- Hover your cursor over the top of largest peak in the MS2 spectrum. What is the mass of this fragment?
- To view a plot that shows more detail, click on the View Vector Metabolomics USI button. Download an image of the tetracycline MS2 plot by clicking on the Download as PNG or Download as SVG.

- 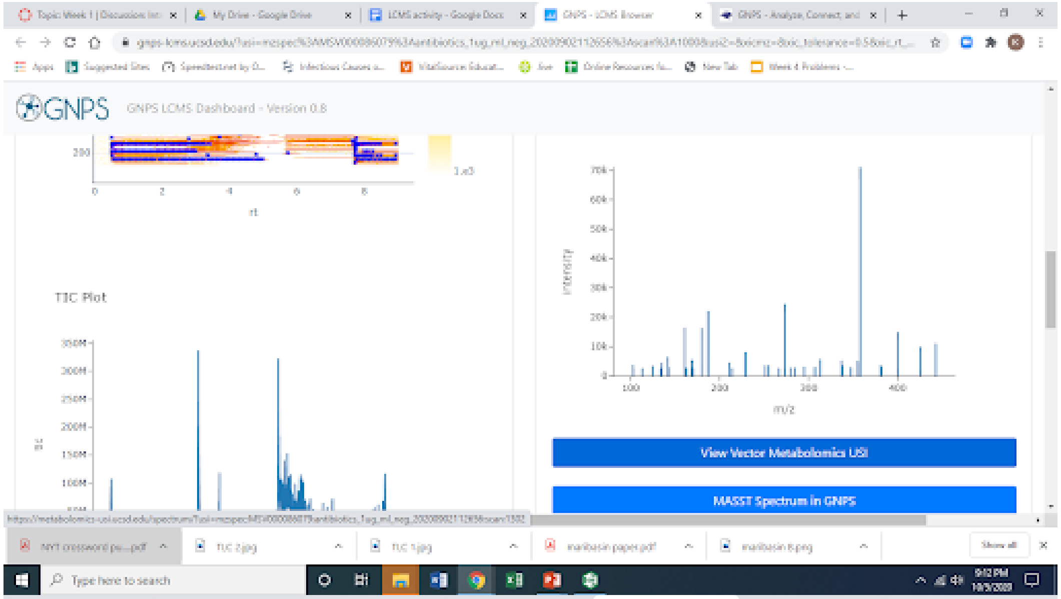


- 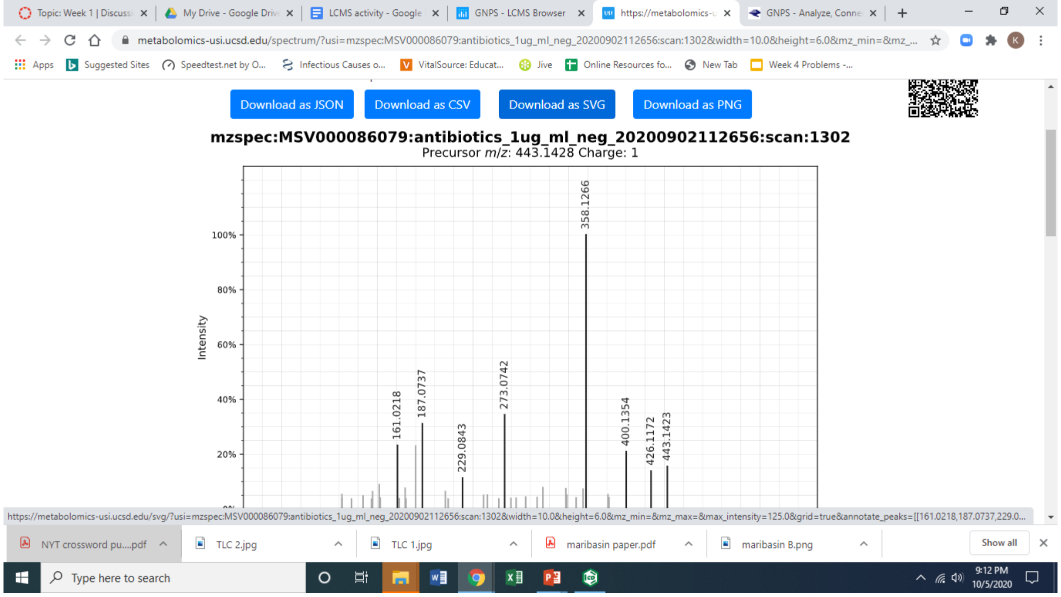


- Below (and linked [here (https://canvas.pointloma.edu/courses/56791/pages/chemdraw-files-for-week-9-lcms)](https://canvas.pointloma.edu/courses/56791/pages/chemdraw-files-for-week-9-lcms) as a ChemDraw file) is a scheme showing some possible fragmentations of tetracycline in negative ion mode, and the resulting fragment ions. Calculate the exact mass for each of these fragment ions. *(Hint: determine the molecular formula of each fragment; note that they* ***already have a negative charge, so you don't need to determine the conjugate base****. Next, calculate the exact mass corresponding to the formula. If you like, you can return to the virtual desktop and use ChemDraw to open the file and determine the formula and exact mass that way!)*
- Using your MS2 data for tetracycline, identify the ion masses in your spectra that could correspond to fragments A, B and C, based on the masses you determined above. Which fragment ion is the most abundant? What does this suggest to you about its stability relative to the other fragments?

- 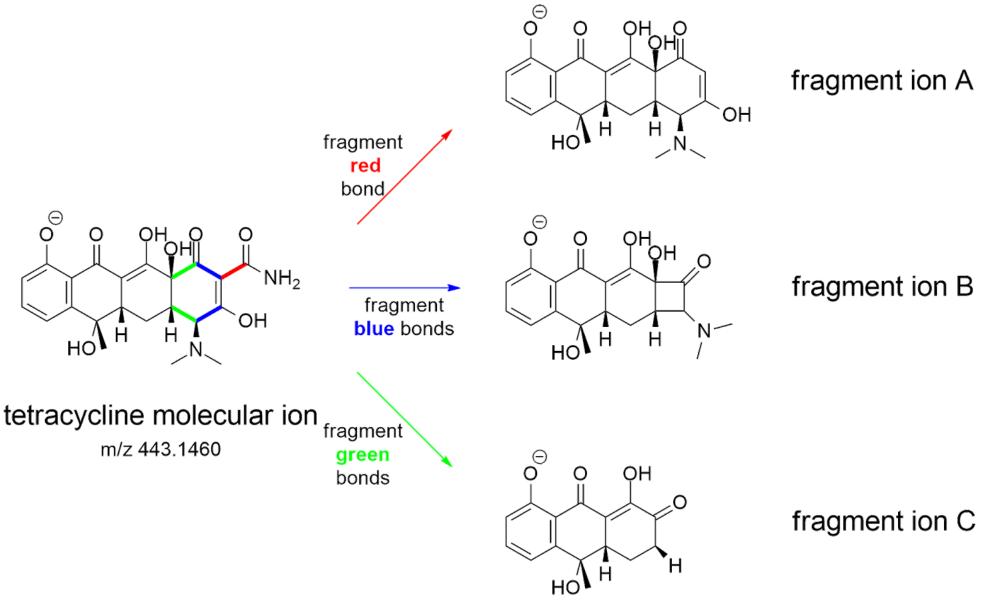


9. Let’s investigate MS2 spectra for two more antibiotics: chloramphenicol and rifamycin.
   - Repeat step 8a-e above to see the MS2 fragmentation spectra for rifamycin and chloramphenicol.
   - The structure of the rifamycin molecular ion is shown below, and in the ChemDraw file **here (https://canvas.pointloma.edu/courses/56791/pages/chemdraw-files-for-week-9-lcms)** , with the most stable fragment shown in blue. In ChemDraw (back on the virtual desktop), use the selection tool to draw a circle around the blue fragment. (Make sure Show Analysis Window is checked under the View menu - and that the Exact Mass is set to display at least 4 significant figures - before selecting the fragment.) What is the exact mass of this fragment? Does this mass match to a peak in the MS2 spectrum for rifamycin?
   - The structure of chloramphenicol is also shown below, and in the attached ChemDraw file. In ChemDraw, use the selection tool to draw circles around sections of chloramphenicol, checking the Exact Mass of each in the Analysis Window. Propose a fragment that might be responsible for the major peak in the MS2 spectrum of chloramphenicol.
   - 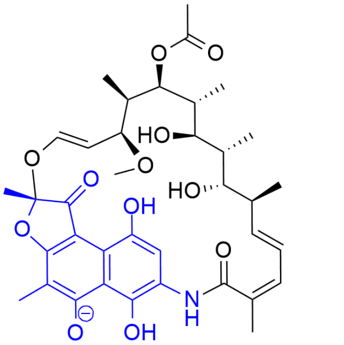

     rifamycin (M-H)$^-$ ion
     probable MS2 fragment shown in blue

     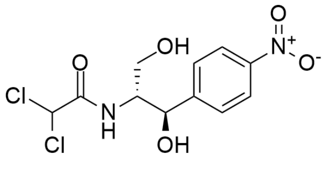

     chloramphenicol

#### Part III: Exploring the LCMS of AH48

For this part, we'll explore data for an extract of AH48 grown at 28 °C in PDB for 5 days - similar to our 'PDB control' extract. Unlike the LCMS data above, this data was collected in **positive ion mode**. As a result, most ions we see will correspond to the molecule plus a proton $(M+H)^+$, also known as the conjugate acid.

1. Return to the [GNPS LCMS Dashboard.](https://gnps-lcms.ucsd.edu/) [(https://gnps-lcms.ucsd.edu/)](https://gnps-lcms.ucsd.edu/)
2. Open the AH48 datafile. Under the **File Selection** tab, in the **GNPS USI** field, delete the default file path and paste **mzspec:MSV000084951:AH48:scan:1000** into the **GNPS USI** box under the **File Selection** tab, then click **Link to these plots**.
3. Let's look at the TIC. Scroll down to the TIC plot. How does it compare to the TIC plot for the antibiotic mixture?
4. Save an image of the TIC plot.
5. Although the TIC for the crude extract is a bit overwhelming, it is possible to use the XIC to 'go fishing' for molecules that might be in there. To see how this can work with AH48, let's try 'fishing' for a certain molecule that I happen to know is in there - we'll call it *Molecule M. Molecule M* has a molecular formula of $C_{49}H_{76}N_{12}O_{14}$. Determine the exact mass for the conjugate acid of *Molecule M*.
6. Paste the exact mass you determined for *Molecule M* into the XIC m/z field to generate an extracted ion chromatogram of only ions of that mass. What do you see? How does this compare to the TIC?
7. Let's look at how we can use a **Library Search** and **MASST Search** to compare our MS2 spectrum to MS2 spectra for known molecules, and to unknown molecules whose MS2 spectra are in the GNPS database. *(This is analogous to the KnownClusterBlast and ClusterBlast searches you did in antiSMASH!)*
    - Click on the little red X over the peak to view the MS2 spectrum corresponding to this peak (compound).
    - 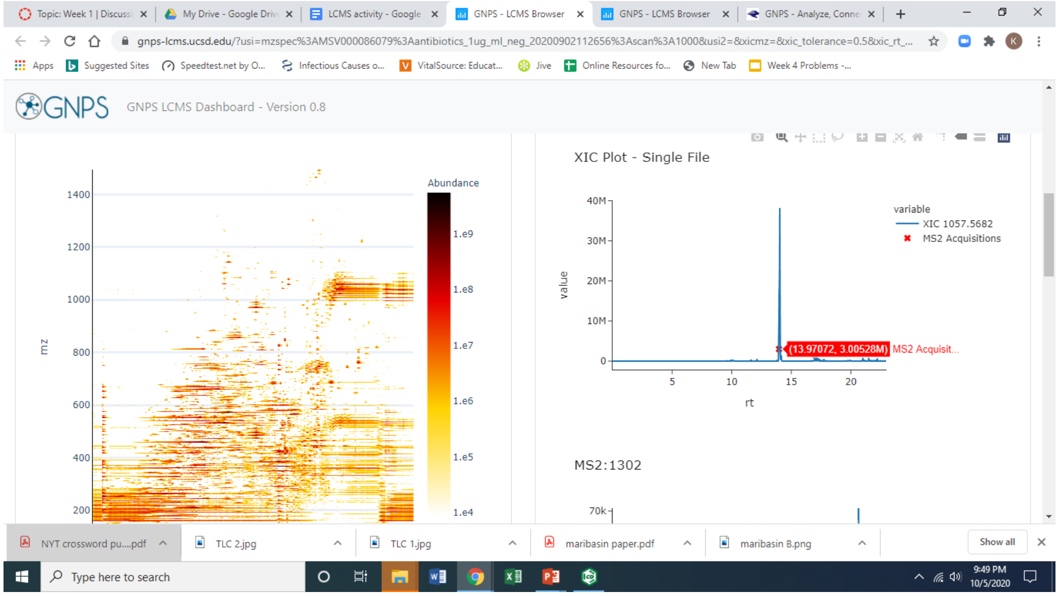

    - Scroll down to the MS2 spectrum. Click on **MASST Spectrum in GNPS**. This will open a new tab with the MASST job page in the GNPS website. (Instead of trying to interpret it ourselves, we're going to use a powerful computer to compare this MS2 spectrum against all the other MS2 spectra in the GNPS database!)

- 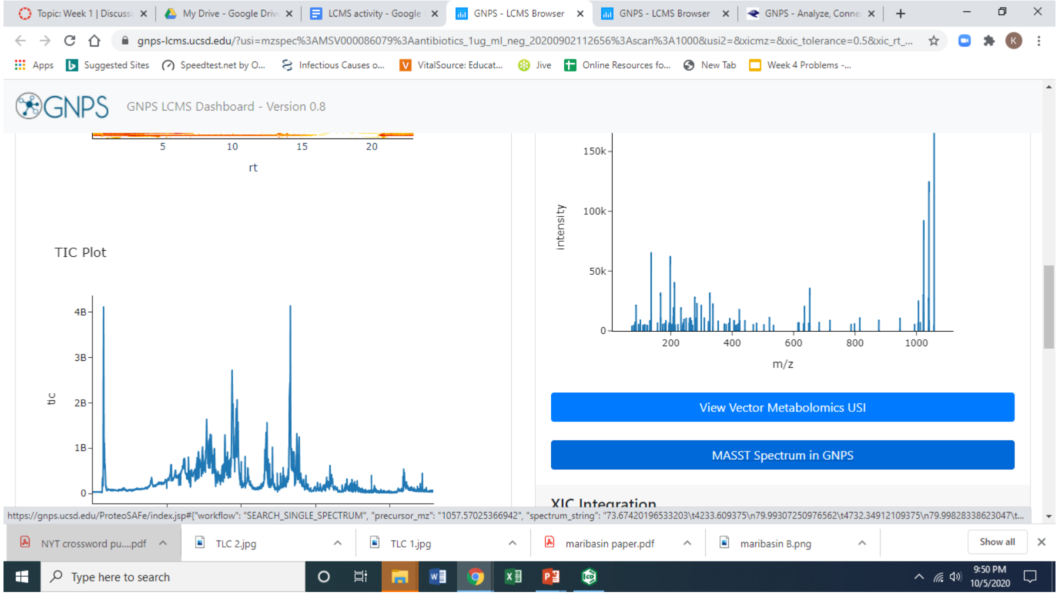
- Give your job a name. Mine was "GNPS MASST search of m/z 1057 MS2 spectrum in AH48 extract".
- 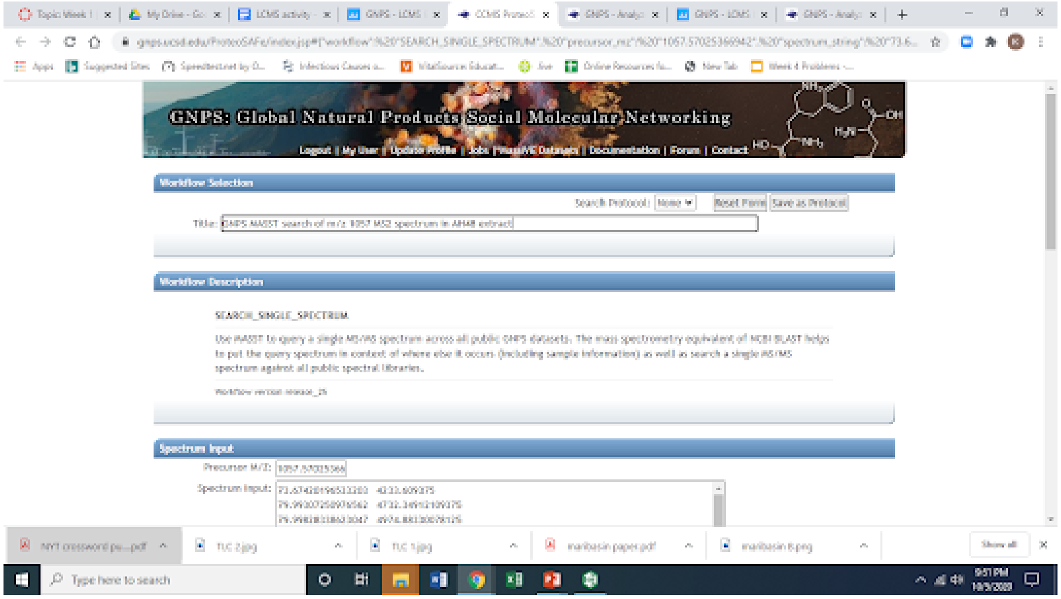
- Verify that your email address appears at the bottom of the page, then click **Submit**. The job will take about 20 minutes to run (during which you can skip ahead to step 8 below; just be sure to come back to finish!)
- When your job is complete, you'll get an email with a link to the completed job page. Open the link to the completed job (or return to the tab if still open).
- At the top, click on **View All Library Hits**. What known molecule does *Molecule M* match to? Scroll all the way to the right to see the structure of this molecule. Right click on the structure to save a copy of the image for your postlab report.

- 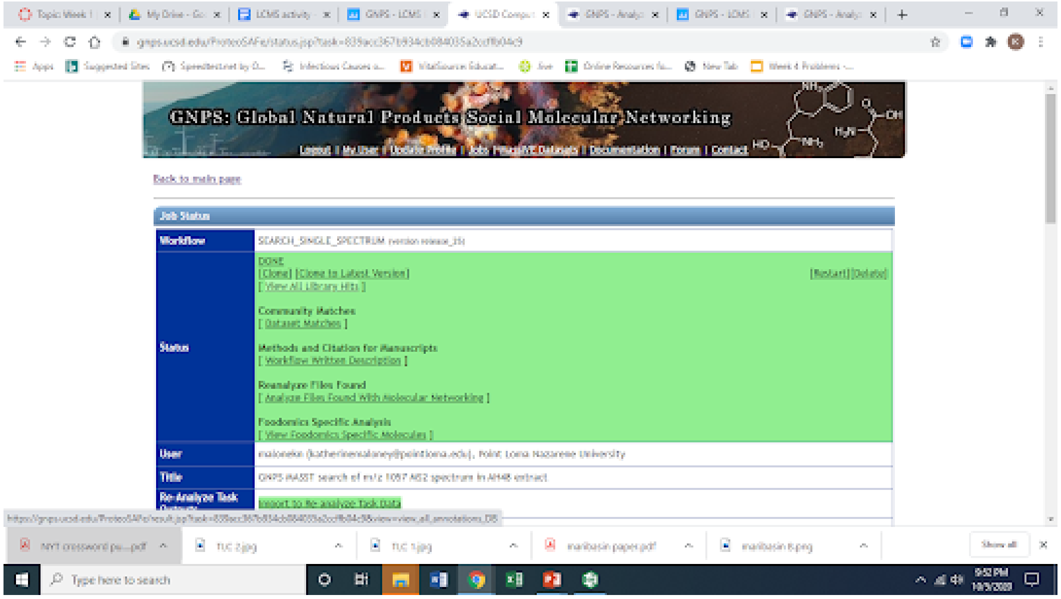


8. Let's go fishing for our Usual Suspects from Week 6 antiSMASH lab.
   - Below is a table of our usual suspects, with the molecular formula given for each. For each, determine the formula and exact mass of the molecular ion peak $(M+H)^+$ (aka, the conjugate *acid*) that you would expect to see in *positive* ion mode.
   - Draw the XIC for each molecular ion determined above. Type the exact mass you determined for each usual suspect in the XIC m/z field at the top right of the page, separating each mass with a semicolon . What do you observe in the XIC plot for each suspect? Do any appear as sharp peaks in the XIC (suggesting there is a molecule with that exact mass in the extracts)? Complete the Usual Suspects table below.
   - Seeing a peak with the same exact mass as one of our usual suspects is good support that there is a molecule with that molecular formula, but it doesn't prove that it is the same molecule. What do we call molecules with the same molecular formula, but whose atoms are connected differently?

**Usual Suspects**

| Compound name | Molecular formula | Conjugate acid formula | Conjugate acid exact mass $(M+H)^+$ | XIC observations |
|---|---|---|---|---|
| butirosin | $C_{21}H_{41}N_5O_{12}$ | | | |
| surfactin | $C_{53}H_{93}N_7O_{13}$ | | | |
| bacilysin | $C_{12}H_{18}N_2O_5$ | | | |
| difficidin | $C_{35}H_{48}O_7$ | | | |
| fengycin | $C_{72}H_{110}N_{12}O_{20}$ | | | |
| bacillaene | $C_{34}H_{48}N_2O_6$ | | | |
| macrolactin H | $C_{22}H_{32}O_5$ | | | |

Congratulations! You've just done a pretty sophisticated analysis of the raw LCMS data from an extract of AH48 - including looking in the data for known molecules (our so-called Usual Suspects). Next time, we'll learn about a technique that lets us extend our analysis in ways that don't require us to have a hypothesis going into it! See you then!
