## Supplemental Information for "GNPS Dashboard: Collaborative Analysis of Mass Spectrometry Data in the Web Browser"

### Use case 1: Mass spectrometry data visualization and linking to molecular networking

**Daniel Petras PhD - Post-Doctoral Scholar - Analytical Chemistry and Chemical Ecology**

*E. coli* Nissle is broadly used as a probiotic and the producer of a series of secondary metabolites including the DNA damaging cytotoxins colibactin and siderophores yersiniabactin. In the context of searching for novel zinc uptake mechanisms, we screened Nissle wildtype and yersiniabactin NRPS knock-out mutants culture extract for yersiniabactin production[1].

The molecular network of MS/MS data and $MS^1$ XICs from the extracts are shown in **SI Figure 1**. Panel a shows the molecular network of yersiniabactins including a series of derivatives around the main compound (m/z 482.124) which are produced by the wildtype but not the NRPS KO mutant according to the network pie charts that visualize MS/MS spectral counts (red = WT, blue = KO). Upon inspection of the extracted ion chromatogram of the m/z 482.124, which is available as a direct link out in the molecular networking results, we could see the abundance difference is evident on the $MS^1$ level in the XIC and the box-plot in panel b. Upon inspection of the XIC and corresponding MS/MS spectra of m/z 482.124, it became apparent that yersiniabactin is present as chromatographically separated isomers, most likely diastereomers with identical MS/MS spectra. Hence classical molecular networking (panel a) that makes use of MS/MS clustering, collapses the spectra to one node[2]. In order to resolve the chromatographically separated features, we reanalyzed the data via feature-based molecular networking using MZmine and GNPS[3], shown in panel c. In the FBMN results, the two yersiniabactin isomers (m/z 482.124) remain as individual nodes in the network. In summary, this is an example of the benefit of direct XIC inspection of classic molecular networking results to verify quantitative results based on MS1 level as well as control of chromatographic peak shape and eventual artifacts such as split peaks or collapsed MS/MS spectra from isomers.

Classic molecular Networking link

GNPS Dashboard for Yersiniabactin link

Feature-based molecular networking link

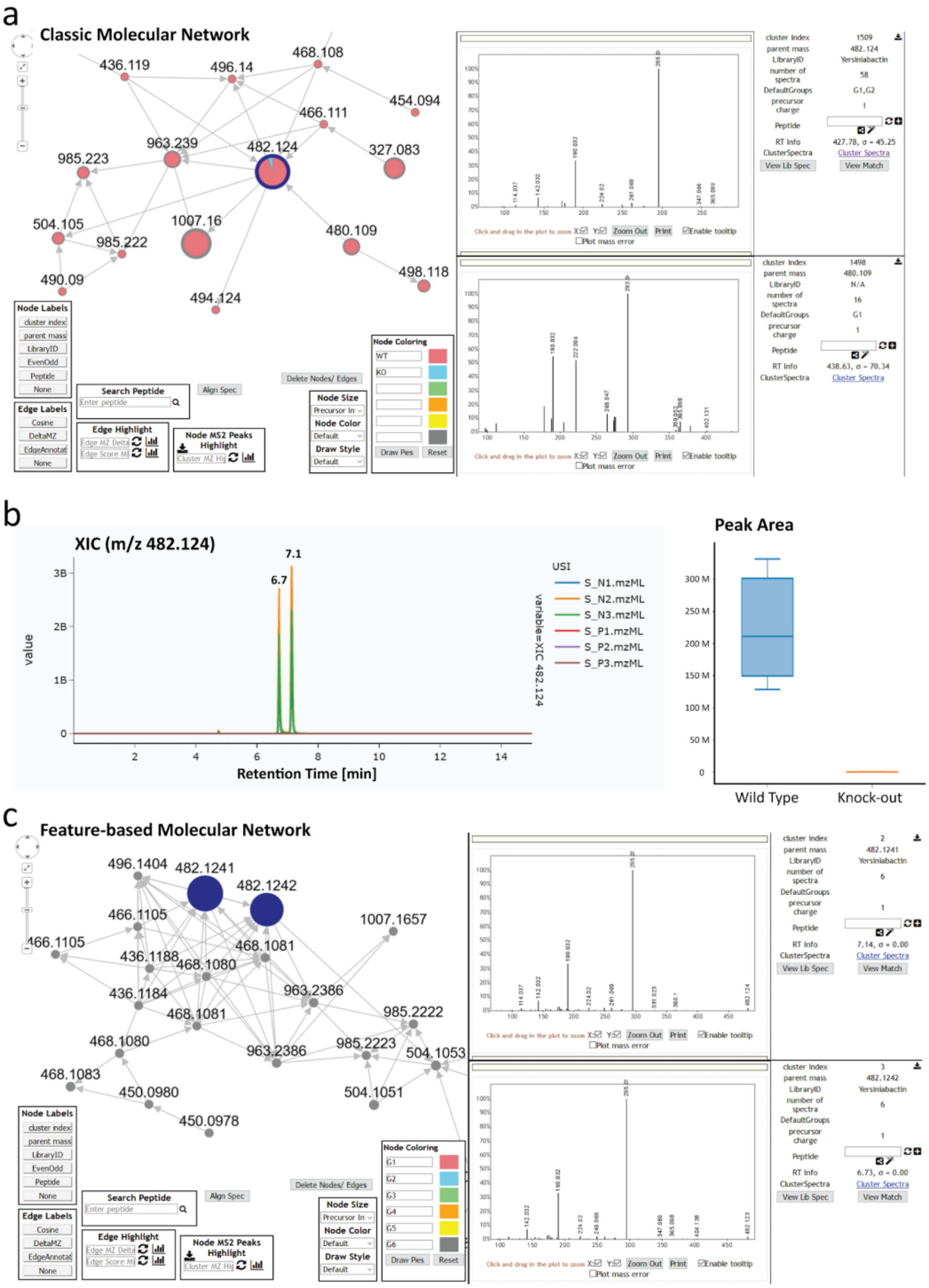


**Supplemental Figure 1: Mass spectrometry data visualization and linking to molecular networks from yersiniabactin from E. coli nissle extracts.** Panel a shows the MS/MS molecular networking results from an *E. coli* nissle wild type and NRPS knock-out strain, indicating the presence of yersiniabactin (m/z 482.124) and its derivatives only in the wild type. Panel b shows the XIC visualization and peak area difference between the WT and KO group as box-plot through the GNPS Dashboard. Panel c shows the feature-based molecular networking results from the same data in which the yersiniabactin isomers (m/z 482.124) are maintained as individual nodes.

### Use case 2: Quick data analysis of MS1 single quad data

**Dale A. Cummings Jr - PhD Student - Biological Chemistry**
**Aaron W. Puri PhD - Assistant Professor - Biological Chemistry**

While many labs now have access to tandem mass spectrometers and can therefore acquire data for MS2-based molecular networking, routine analysis is often still performed on unit resolution MS1 mass analyzers, such as single quadrupoles. However, proprietary software often makes it difficult to rapidly analyze and share these types of data. To address this we have added functionality for analyzing MS1 spectra in the GNPS Dashboard specifically in netCDF format, which is commonly used by single quadrupole LC-MS systems.

As an example use case, we analyzed the metabolome of the methylotrophic organism *Methylorubrum extorquens* PA1 (PA1) using an Agilent 6120 single quadrupole LC-MS. Methylotrophs have previously been reported to biosynthesize different natural products when grown on single versus multicarbon substrates[4,5]. We grew PA1 in minimal media containing either methanol or succinate as the sole source of carbon and energy, and subsequently extracted the supernatant with acidified ethyl acetate. We then analyzed this extract by LC-MS and examined the results using the GNPS Dashboard.

One difference in the metabolome under these two growth conditions appears to be the production of quorum sensing signals. PA1 possesses a single LuxI-family acyl-homoserine lactone (acyl-HSL) synthase with 100% amino acid identity to MlaI[4]. In the closely related strain *Methylorubrum extorquens* AM1 (AM1), Vorholt and coworkers demonstrated that MlaI produces the long-chain quorum-sensing signals 7*Z*-$C_{14}$-HSL and 2*E*,7*Z*-$C_{14}$-HSL when grown on methanol but not succinate[5]. We used the GNPS Dashboard to generate XICs for the protonated masses of these signals (*m/z* 308.1 for 2*E*,7*Z*-$C_{14}$-HSL, and *m/z* 310.1 for 7*Z*-$C_{14}$-HSL) (Figure S2a). These masses are apparent at retention times of 25.9 minutes for *m/z* 308 and 26.3 minutes for *m/z* 310 in the methanol-grown culture, but not in the succinate culture, which can be quantified by comparing the automatically generated peak integrations in the two conditions (Figure S2b). These features therefore likely correspond to quorum-sensing signals 7Z-C14-HSL and 2E,7Z-HSL, and demonstrate how the GNPS Dashboard can be used to rapidly analyze MS1 data.

GNPS Dashboard link.

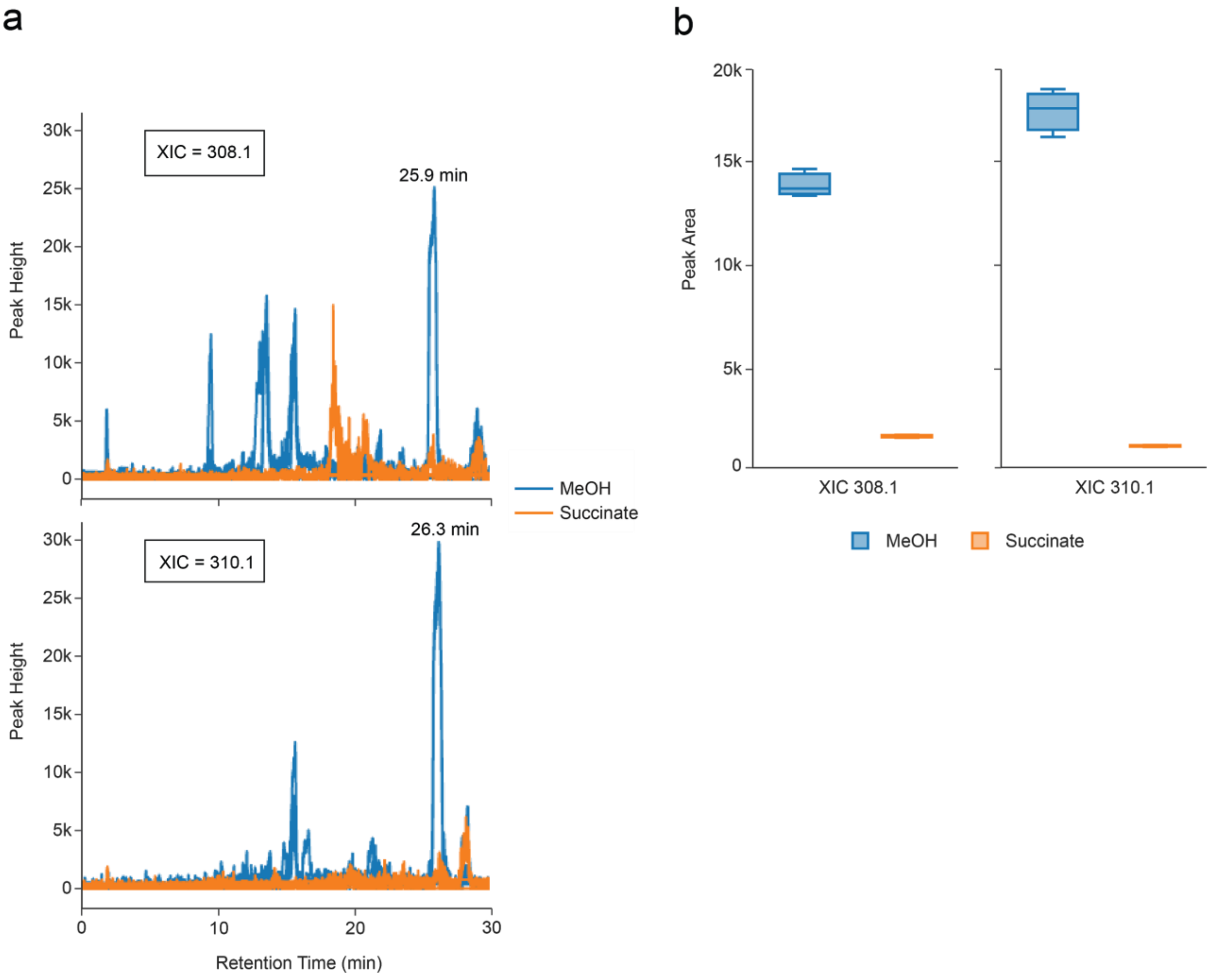


**Supplemental Figure 2: Quick data analysis of MS1 single quad data**
Identification of differences in the extracellular metabolomes of the methylotrophic bacterium *Methylorubrum extorquens* PA1 (PA1) grown on different carbon and energy sources. (a) XICs for protonated masses corresponding to quorum-sensing signals produced by PA1. (b) Box plots of integrated peak areas for the same XICs from a retention time window of 25-27 minutes. Data are from three separate cultures for each growth condition.

**Use case 3: Collaborative Exploration of LC-MS Data**

**Mingxun Wang PhD - Post-Doctoral Scholar - Computer Science and Bioinformatics**
**Daniel Petras PhD - Post-Doctoral Scholar - Analytical Chemistry and Chemical Ecology**

The exploration of non-targeted mass spectrometry is often a long and collaborative process, e.g. between collaborating researchers or between mentors and mentees. In many instances, it is difficult to collaboratively explore and share results when it is impractical or impossible to physically sit together, for example during a global pandemic. With the GNPS Dashboard, it is possible to share exactly the settings and visualization that a researcher is seeing on their own screen either by our collaborative sync feature or by simply copying and pasting a url. This sharing ability enables others to build upon this visualization and reshare to collaboratively enhance and dive deeper into the data, whether in joint data analysis sessions with collaborators, reports, or peer-reviewed publications. One particular example and collaborative visualization dialog can be seen below with quality control (QC) sample of a mixture of 6 standard compounds (**SI Figure 3**):

Researcher A starts by initial data loading of the QC data- link.

Next, Researcher B has the information of exact masses and adds XIC for the 6 standard compound used (m/z 271.0315; 278.1902; 279.0909; 285.0205; 311.0805; 314.1381) - link.

To verify the ID of one of the standards through its isotope pattern, Researcher A then visualizes of MS1 spectrum for the m/z 314.1381 standard - link.

Finally, Researcher B inspects co-eluting features by zooming in on 2D LC-MS heatmap for elution of the m/z 314.1381 standard - link.

The final url can then be shared with others in the team and used to protocol the instrument performance for this particular experiment.

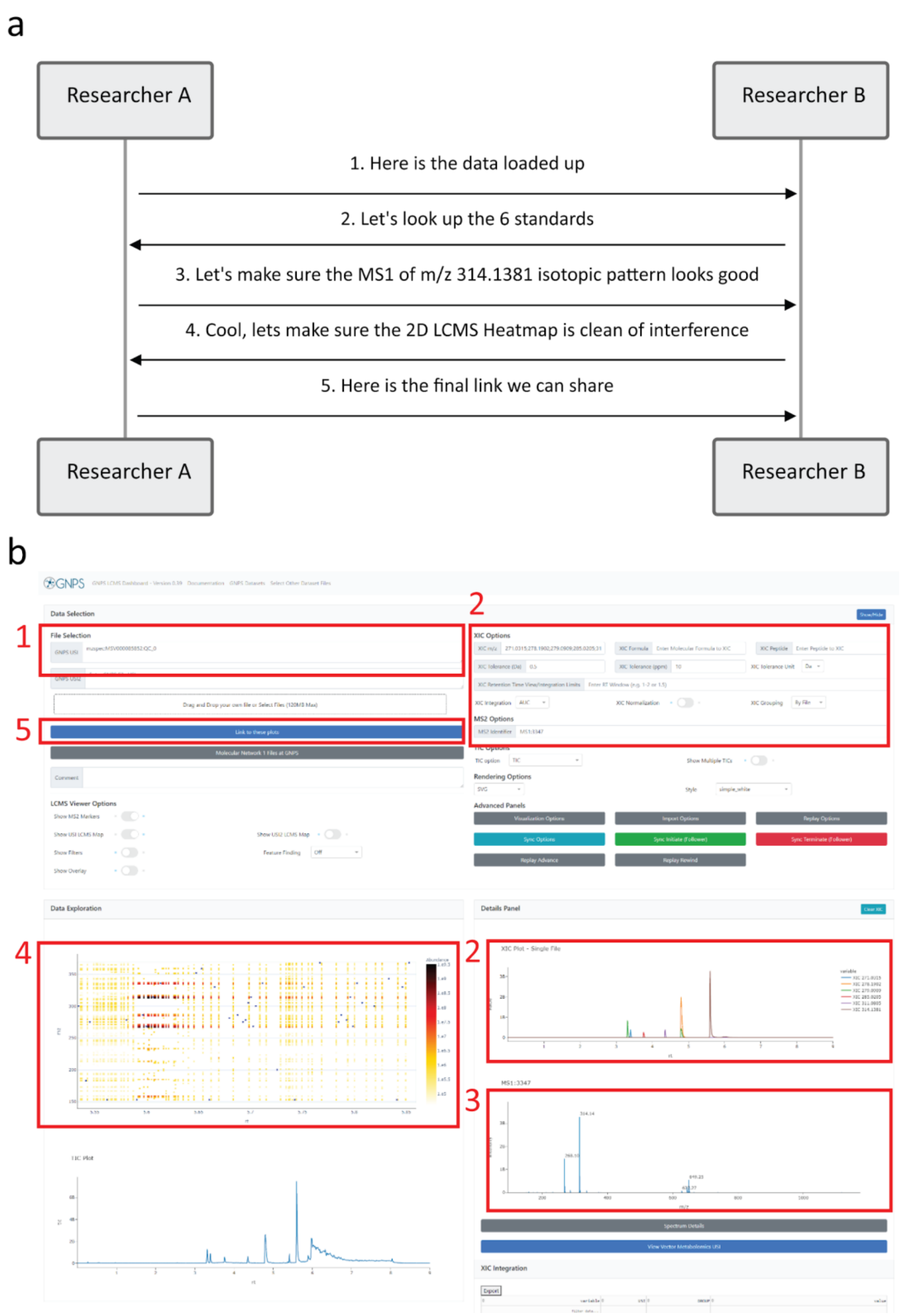


**Supplemental Figure 3:** In panel a we see an example communication between two users to collaboratively analyze a mass spectrometry sample. This communication can be done purely remotely by sharing URLs or the synchronization feature in GNPS Dashboard. In panel b we can see how the discrete steps line up with updating different parts of the dashboard as the analysis progresses.

### Use case 4: Inspection and validation of published results based upon public data sharing/inspection

**Mingxun Wang PhD - Post-Doctoral Scholar - Computer Science and Bioinformatics**
**Daniel Petras PhD - Post-Doctoral Scholar - Analytical Chemistry and Chemical Ecology**

Data transparency and verification of software-derived results have become key aspects in recent years in proteomics and metabolomics publications thanks to policy and cultural shifts within these fields. It has even become the standard for many scientific communities where publishers require data sharing for papers to be accepted for publication. While there are differences in guidelines with regards to data content (e.g. raw and processed data), as well as accepted repositories (e.g. ProteoXchange[6] initiative or Nature Publishing Group guidelines), there still exist multiple barriers for reviewers and readers of scientific articles to make use of the shared data for inspection or reproduction of the published results.

The GNPS Dashboard in combination with the GNPS Dataset Explorer enables direct visualization of public data from the most common mass spec data repositories (PRIDE[7], MassIVE[8]/GNPS[2], MetaboLights[9], Metabolomics Workbench[10]) in a vendor-independent fashion. The GNPS Dashboard removes the need to install software and to download MS data locally and enables inspection of and reproduction of peak visualization and integration as well as downstream data analysis in the GNPS environment within a few clicks in the web browser.

Between October 2020 and January 2021, we inspected 10 LC-MS datasets from peer-reviewed articles as well as preprints in the fields of natural product research, metabolomics, and proteomics that were shared through the social media platform Twitter, which is quickly becoming a fundamental part of scientific literature sharing and discussions. In all cases, we inspected the raw MS/MS data using the GNPS Dashboard and created extracted ion chromatograms (XIC) for the masses of the main compounds discussed in the manuscripts. Depending on the experimental design of each study we also reproduced simple quantitative analysis by integrating the XICs and comparing the peak areas of the experimental groups as box plots directly in the GNPS Dashboard. Upon positive validation, we provided direct feedback through the comment and retweet function in the social media platform, including confirmation of the presence of described masses, MS/MS matching, and quantitative differences between sample groups (**SI Figure 4**), as well as comparison to other public data through repository scale analysis (**SI Figure 5**), and recommendations for additional data analysis options such as molecular networking in the case of one of the preprints (**SI Figure 6**).

#### Example Tweets List

1. Tweet - Dashboard - Comparative Metabologenomics Analysis of Polar Actinomycetes
2. Tweet - Dashboard - Exploring the Chemical Space of Macro- and Micro-Algae Using Comparative Metabolomics
3. Tweet - Dashboard - Species Prioritization Based on Spectral Dissimilarity: A Case Study of Polyporoid Fungal Species

4. Tweet - Dashboard - Planomonospora: A Metabolomics Perspective on an Underexplored Actinobacteria Genus
5. Tweet - Dashboard - AdipoAtlas: A Reference Lipidome for Human White Adipose Tissue
6. Tweet - Dashboard - Convergent evolution of pain-inducing defensive venom components in spitting cobras
7. Tweet - Dashboard - Iron-meditated fungal starvation by lupine rhizosphere-associated and extremotolerant Streptomyces sp. S29 desferrioxamine production
8. Tweet - Dashboard - Bacterial–fungal interactions revealed by genome-wide analysis of bacterial mutant fitness
9. Tweet - Dashboard - Deep Interrogation of Metabolism Using a Pathway-Targeted Click-Chemistry Approach
10. Tweet - Dashboard - MassIVE.quant: a community resource of quantitative mass spectrometry-based proteomics datasets

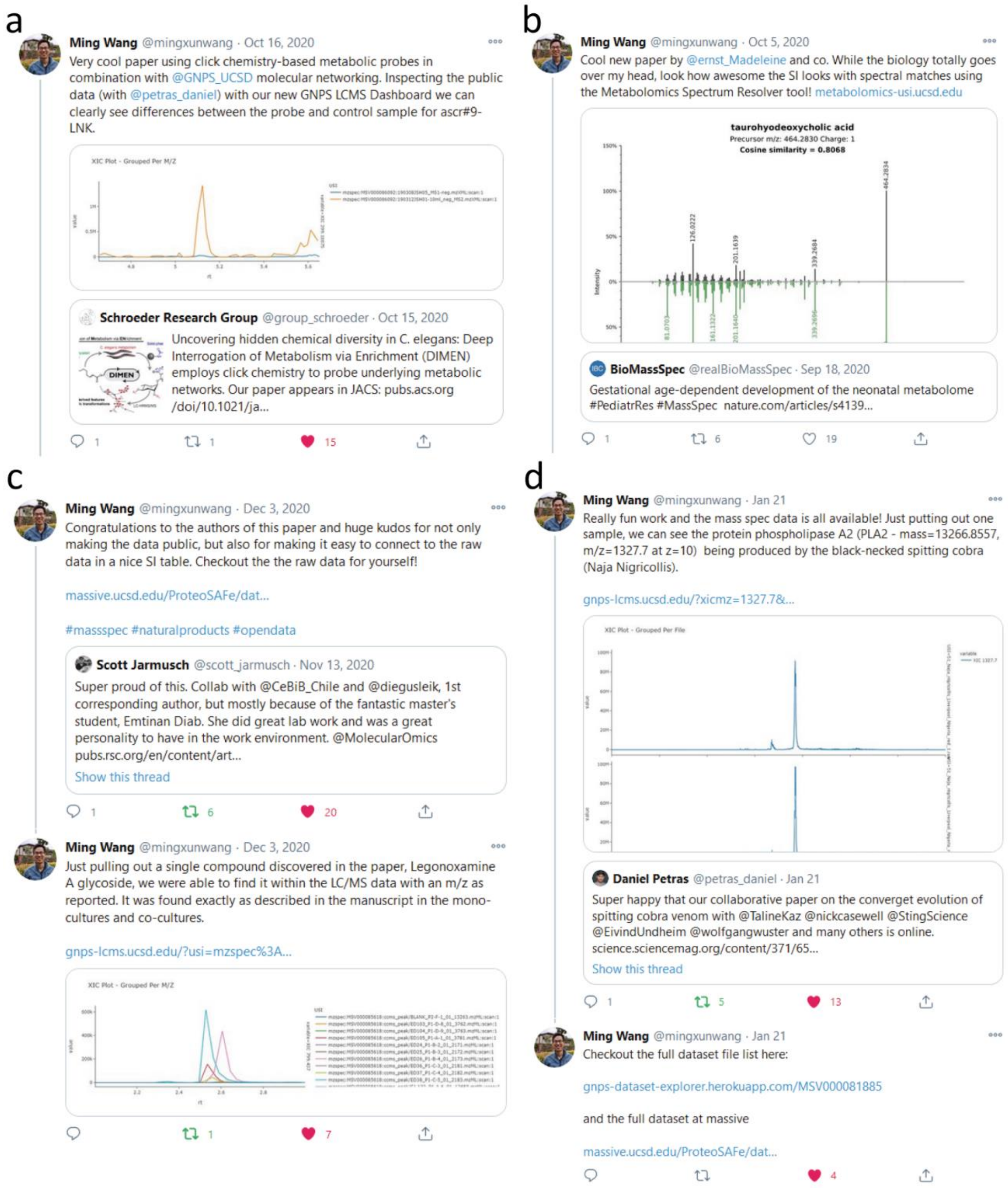


**Supplemental Figure 4 -** Inspection of publicly shared mass spectrometry data from Deep Interrogation of Metabolism Using a Pathway-Targeted Click-Chemistry Approach

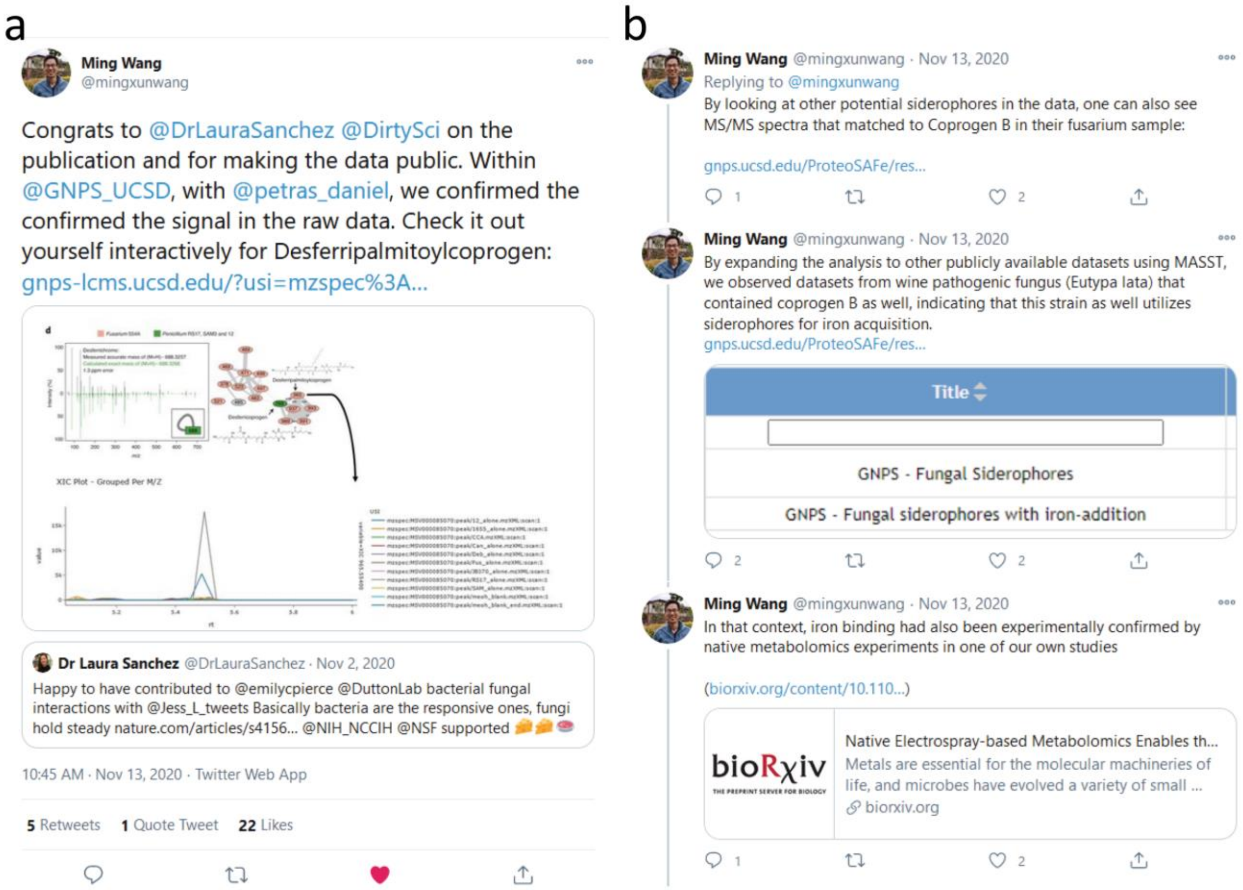


**Supplemental Figure 5:** Inspection of publicly shared mass spectrometry data from Bacterial–fungal interactions revealed by genome-wide analysis of bacterial mutant fitness

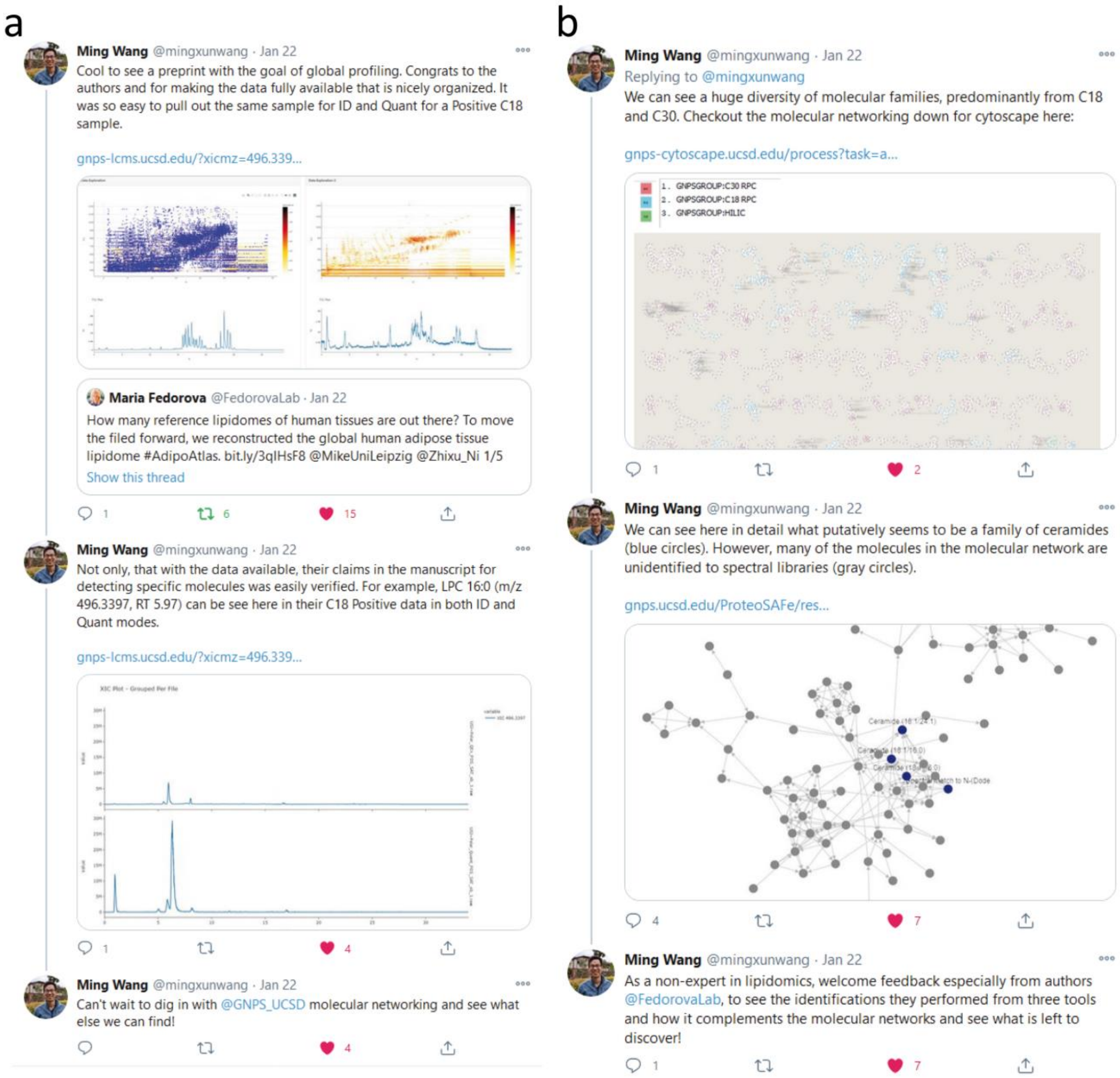


**Supplemental Figure 6:** Inspection of publicly shared mass spectrometry data of AdipoAtlas: A Reference Lipidome for Human White Adipose Tissue

**Use Case 5A: GNPS Dashboard usage in Teaching Setting**

**Katherine Maloney PhD - Professor - Natural Product Chemistry,**
**Allegra Aron PhD - Post-Doctoral Scholar - Biological Chemistry**

GNPS Dashboard proved useful in a teaching setting, as we developed a virtual, interactive laboratory exercise for undergraduate students to explore LC-MS data generated by the instructor, as part of Tiny Earth, an initiative to crowdsource antibiotic discovery in a Course-based Undergraduate Research Experience (CURE) GNPS Dashboard was an ideal tool for this interactive (or a virtually interactive) lab because software installation was not necessary and software licenses did not need to be obtained, both challenging obstacles when working with a class. One option frequently employed is the use of a virtual desktop that has the requisite software installed. For example, we used such a system for the processing of NMR data in the Tiny Earth[11] course. However, students found the virtual desktop to be clunky and error-prone. The web-based interface of the GNPS Dashboard proved to be markedly more user-friendly, giving reproducible and shareable results. Moreover, the instructor can equip students with path links to data for exploration in GNPS Dashboard or can take students through data exploration using the synchronization feature.

The interactive lab exercise making use of the GNPS Dashboard was part of the Tiny Earth Chemistry pilot course (CHEM 1003L) developed and taught by Katherine Maloney and Allegra Aron at Point Loma Nazarene University in San Diego, CA. The goal of the exercise was to teach students about LC-MS experiments and data structure by providing hands-on experience with LC-MS data generated from pure antibiotic standards and antibiotics in culture extracts. Briefly, the exercise asks students to determine the exact mass of antibiotics in question; once this has been completed students explore the LC-MS of a mixture of antibiotics using the LC-MS Viewer and a path link provided in the lab protocol. Students explore the TIC plot to load the MS1 spectrum for each antibiotic (**SI Figure 7**). Next, students explore the XIC for each antibiotic then hone in to inspect the tandem MS (or MS2) spectrum of each antibiotic, trying to annotate fragment peaks with particular molecular fragments. Finally, students are given another path link for the LC-MS of culture extracts. Students are tasked with first looking through the TIC and XIC plots; from here, students dive into further exploration of compounds of interest in GNPS, including the use of Mass Spectrometry Search Tool (MASST)[12]. The full laboratory exercise is provided as **SI Data 1**.

GNPS Dashboard - Link

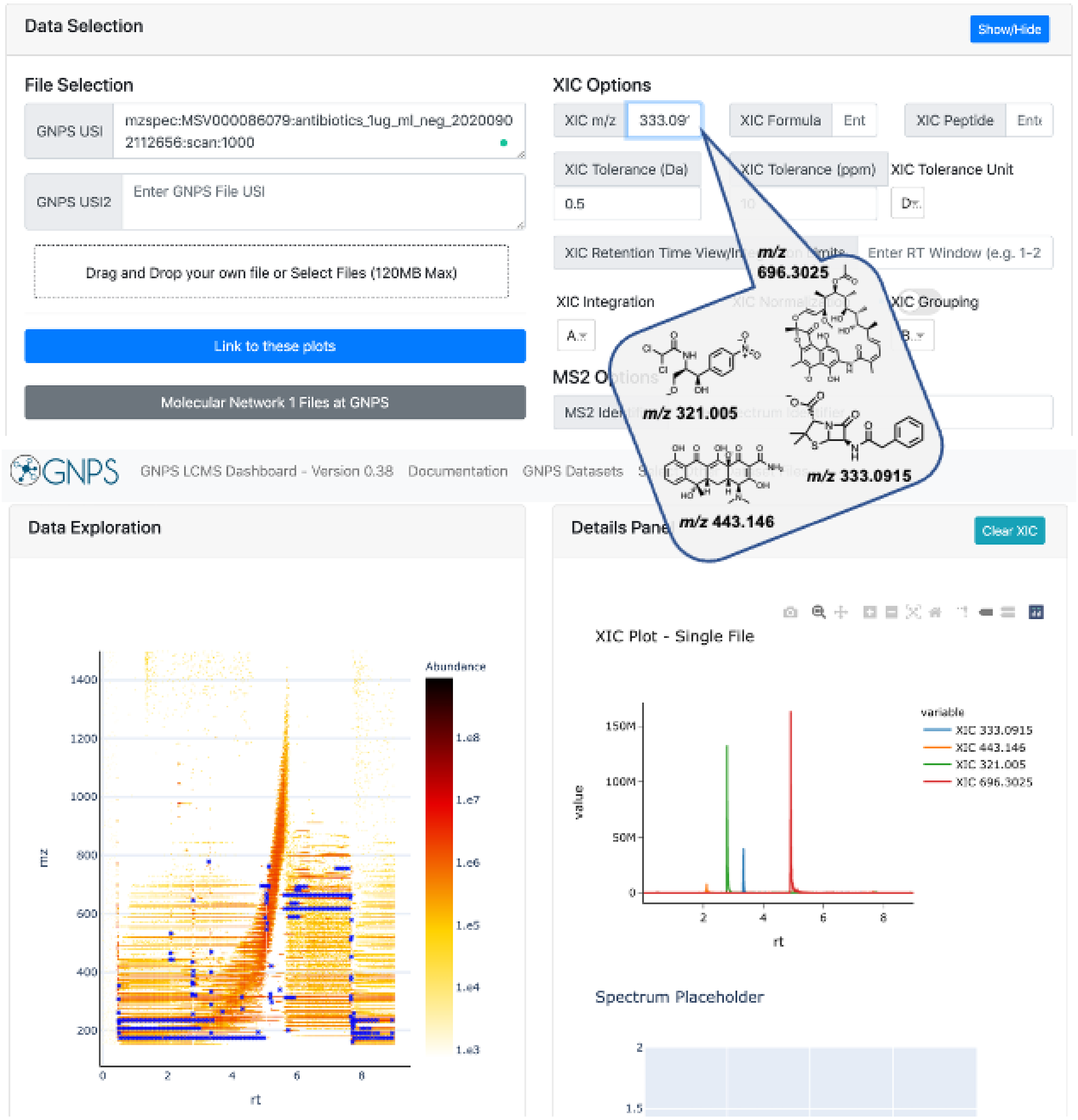


**Supplemental Figure 7:** Students use the GNPS Dashboard (link) to visualize XIC plots of antibiotics of interest, which include chloramphenicol ([H-H+], *m/z* 321.005), tetracycline ([H-H+], *m/z* 443.146), penicillin G ([H-H+], *m/z* 333.0915), and rifamycin ([H-H+], *m/z* 696.3025).

**Use Case 5B: GNPS Dashboard for Targeted MS in a Teaching Setting**

**Michael Marty PhD – Assistant Professor – Analytical Chemistry**
**Deirdre Belle-Oudry PhD – Senior Lecturer – Analytical Chemistry**

In addition to analyzing LC-MS data generated by the instructor or in public data sets, the GNPS Dashboard is also useful for undergraduate students analyzing targeted data for quantitative analysis. For example, the GNPS Dashboard was used as the data analysis tool for two new labs in the *Quantitative Analysis Lab* (CHEM 326) course at the University of Arizona in Tucson, AZ developed by Michael Marty and Deirdre Belle-Oudry. Earlier in the semester, the students used HPLC with UV detection to quantify aspirin, acetaminophen, and caffeine[13]. However, additional peaks can be present in the UV chromatogram due to hydrolysis of acetyl-salicylic acid into salicylic acid. In the first LC/MS lab, students use LC/MS/MS with a triple quadrupole mass spectrometer to first identify the degraded product from product ion scans and then use single reaction monitoring (SRM) scans for targeted quantitation. After collecting the data and converting it to mzML format using MSConvert[14] students use the GNPS Dashboard to view the MS1 and MS2 data to assign the peaks. Example data is posted to MassIVE MSV000087058 to allow students to access training data from a direct link. Next, they use the Dashboard to view and integrate the SRM chromatograms. Because the LC/MS/MS system has a UV detector, students can also load the UV chromatograms into the GNPS Dashboard to compare retention times from both detection methods. In the second lab, the students apply the skills they develop in the first lab to quantify amino acids in food samples based on targeted SRM scans. Overall, the GNPS Dashboard provides a fast and user-friendly method to quantify targeted SRM scans and view MS data. The web-based interface provides easy and inclusive access for students with all different types of computers.

Tutorial Video - https://www.youtube.com/watch?v=tT5b1IIJCE8

GNPS Dashboard - Link

**Use Case 6: Data Visualization for Quantitative Proteomics Results**

**Ben Pullman - PhD Student - Computational Mass Spectrometry**
**Nuno Bandeira PhD - Professor - Computational Mass Spectrometry**

MassIVE.quant[15] provides a platform for the sharing of quantitative re-analyses, from RAW data to features to differential expression (DE) analysis. The complete set of scripts and commands used to run the experiment enables reproducible data reanalysis and also facilitates altering parameters and algorithms used for slightly-modified future reanalyses. Incorporating the GNPS Dashboard provides a more interactive exploration of these stages of analysis and further facilitates investigating reported findings.

In the IPRG2015[16] study (MSV000079843/PXD015300), four samples are spiked into a consistent yeast background in six known, different concentrations. There are four conditions, each with a different known combination of the previous four spikes in samples (C1-C4), and each condition is run three times as technical replicates. The experiment design proves itself useful when considering quantifications pipelines, as we can assess if the differential expression found matches what is expected. In the reanalysis: RMSV000000249.18, which uses MaxQuant[17] for feature finding, almost all true positives (TP) are found, and the number of false positives (FP) is overall very low. However, there are still some FPs to investigate. sp|P07834|CDC4_YEAST is not a spiked in protein but is DE between two conditions in the experiment, C4 as compared to C2, and this expression is supported by only a single feature, though the feature does have consistent intensity across all three replicates in the sample (**SI Figure 8a**, Protein Differential Expression Results). We can find the location of the features supporting the peptide by looking in the MaxQuant[17] analysis (**SI Figure 8b**, Peptide Differential Expression Results).

To further examine the connection between the differential results and the raw data, the GNPS Dashboard enables us to examine the region where this peptide was detected to be, and inspect if there might be any other artifacts that might be influencing this feature intensity. We see that there's a feature for a different protein group that appears in exactly the same area and the first isotope of that feature overlaps completely with the monoisotopic *m/z* of the sp|P07834|CDC4_YEAST feature. Also generally it seems that the region around where the feature was acquired has many features (**SI Figure 8c**, GNPS Dashboard Link).

Finally, to confirm, when looking at the MS2 mapped to the feature, we also see that most of the intensity for the MS2 being considered cannot be explained by the peptide which it is assigned, lending more evidence to the fact that this feature might be improperly considered. (SI Figure 8d, Peptide Fragmentation Visualization - mzspec:PXD015300:JD_06232014_sample4_C:scan:8576:LSQKYPK/2).

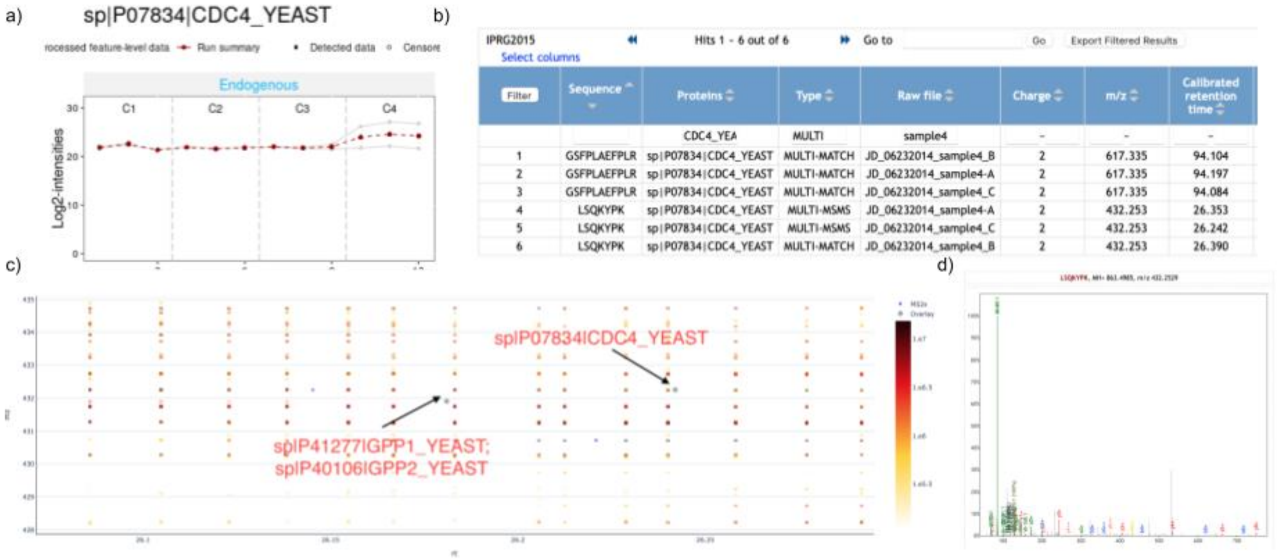


**Supplementary Figure 8: Connecting Protein Differential Expression with Raw Proteomics Data -** a) shows the log2 intensity for features that map to sp|P07834|CDC4_YEAST in all 4 conditions from the IPRG2015 experiment, illustrating how the intensity is consistent across conditions 1-3 but significantly upregulated in condition 4, though only due to one features abundance, b) shows all features in Condition 4 that are for this protein, 6 in total, corresponding to 2 peptides, each with 3 replicates c) shows the LC-MS map for the region near the feature for LSQKYPK, the feature with very high abundance, in the 3rd replicate, and d) shows the annotated MS/MS spectrum for the peptide, and how the annotated peaks don’t show much of the intensity of the spectrum.

### Use case 7: Data visualization and parameter optimization for large scale feature finding

**Carlos Molina-Santiago PhD, Post-Doctoral Scholar, Microbiology**

*Bacillus subtilis* is a well-known Gram-positive strain that exhibits great potential as a biocontrol agent against bacterial and fungal pathogens in plants, such as in cucurbit crops[18]. In the example, we showcase the study of the metabolic changes occurring on melon leaves after the inoculation of *B. subtilis* by non-targeted LC-MS/MS. We compared the metabolomes of untreated and treated melon leaves 5 days after inoculation with *B. subtilis*.

The optimization of feature finding parameters is one of the most important steps in the analysis of non-targeted LC-MS data, as it dictates much of the downstream quality of results. The GNPS Dashboard enables the tweaking of feature-finding parameters and immediate visualization of them for further downstream large-scale feature-finding via MZmine[19] as a GNPS workflow.

We have used the GNPS Dashboard as a fast and easy tool to firstly compare the samples. We initially checked the successful colonization of the melon plant by *B. subtilis* by displaying the presence of surfactin (*m/z* 1036.69), a secondary metabolite known to be produced by this bacterium[20,21]. The GNPS Dashboard (**SI Fig 9A and B**) confirming its presence and abundance in treated leaves. With all the data already loaded in the GNPS dashboard, we can now perform Feature Finding directly modifying the settings in this tool (**SI Fig. 9C**) obtaining a heatmap where we can identify all the features detected (green diamonds) and their intensity coverage (Fig. XD). Once confirmed that the data are of enough quality to perform a deepest analysis, we can now perform Feature-Based Molecular Networking (FBMN)[3] and MZmine to all the datasets directly from the GNPS dashboard. With that, we can now compare all the samples of the dataset using PcoA analysis (**SI Fig. 9E**) confirming the pattern differences between non-treated and treated melon leaves and visualizing the results using Cytoscape. These tools have permitted us, for instance, to highlight the differences observed in a cluster where we can find known potential antimicrobial compounds produced by plants such as 3,5-Dimethoxy-4-hydroxycinnamic acid, neochlorogenic acid, and trans-ferulic acid (**SI Fig. 9F**) In this case, we find a higher abundance of these compounds when melon leaves are treated with *B. subtilis*, a result that we can also confirm using box plots from the GNPS dashboard (**SI Fig. 9G**), showcasing the seamless interlinking of the GNPS Dashboard to optimize and launch data analysis as well as downstream manual inspection and confirmation of results from FBMN[3].

GNPS Dashboard link

FBMN GNPS Task link

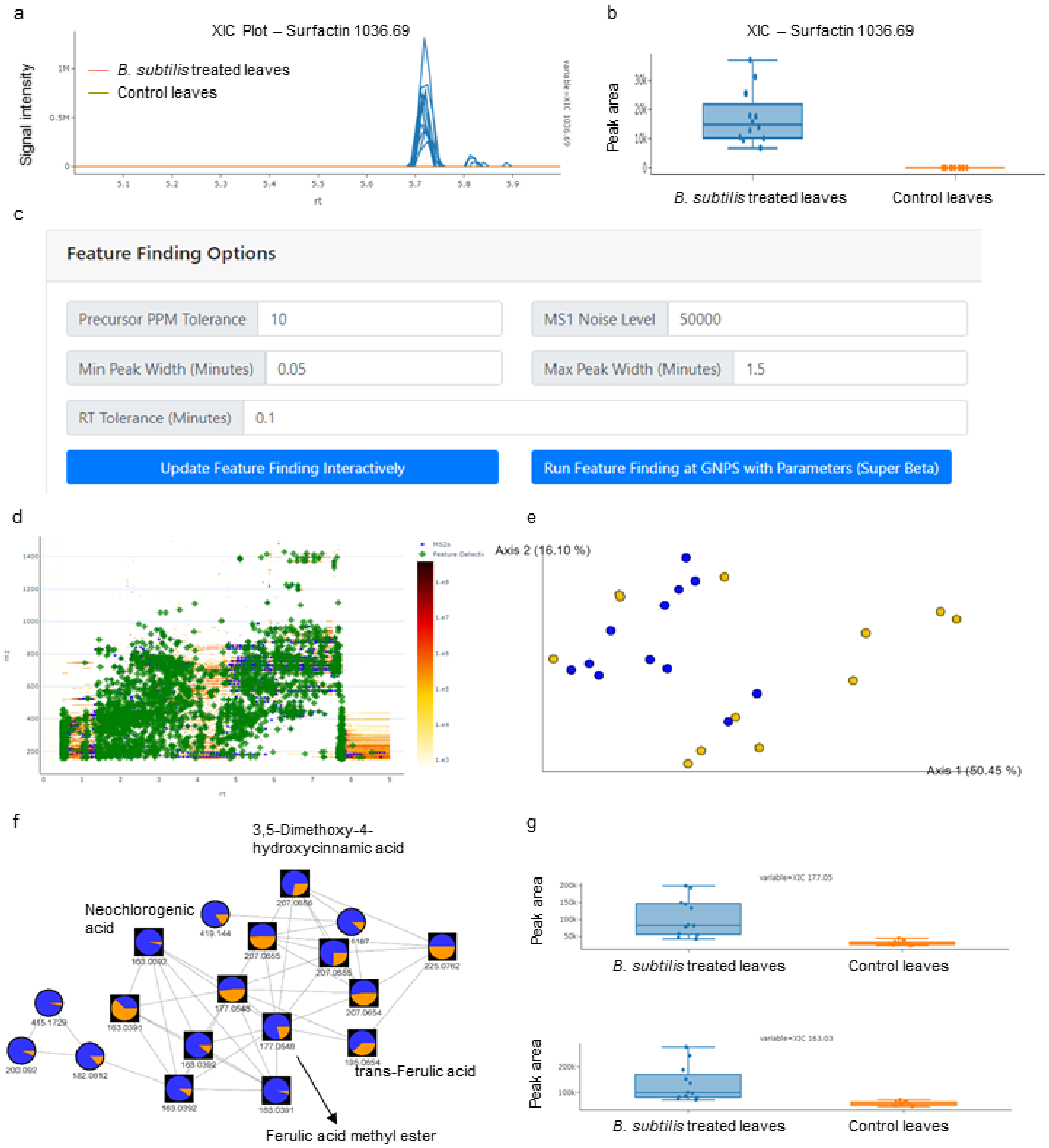


**Supplemental Figure 9. Optimization of feature finding settings and feature-based molecular networking through the GNPS dashboard.** Panel a shows the comparison of XIC of Surfactin (*m/z* 1036.69) of B. subtilis treated plants (blue) and control plants without treatment (orange) in the GNPS dashboard. Box plots of integrated peak areas of Surfactin of treatment vs. control are shown in panel b and show that surfactin was detected in the treated sample but not the control. In the GNPS Dashboard MZmine parameters were visually optimized and shown as green boxes in the 2D LC-MS heatmap in panel d for a representative sample and then directly applied in a link out for feature finding using the GNPS MZmine workflow. Panel e shows the multivariate (PCoA)[22] visualization of sample-to-sample distance based on all features in the data set. Panel f shows a selected network family that show different abundances between treatment and control group which was then further inspected by manually integrating the XICs of ferulic acid methyl ester (*m/z* 177) and neochlorogenic acid (m/z 163) in the GNPS Dashboard as shown in panel g.

**Use case 8: Outliers in data analysis - Visualization of LC-MS can diagnose upstream data issues**

**Rachel Neve - PhD Student - Microbiology**
**Vanessa Phelan PhD - Assistant Professor - Chemical Ecology**

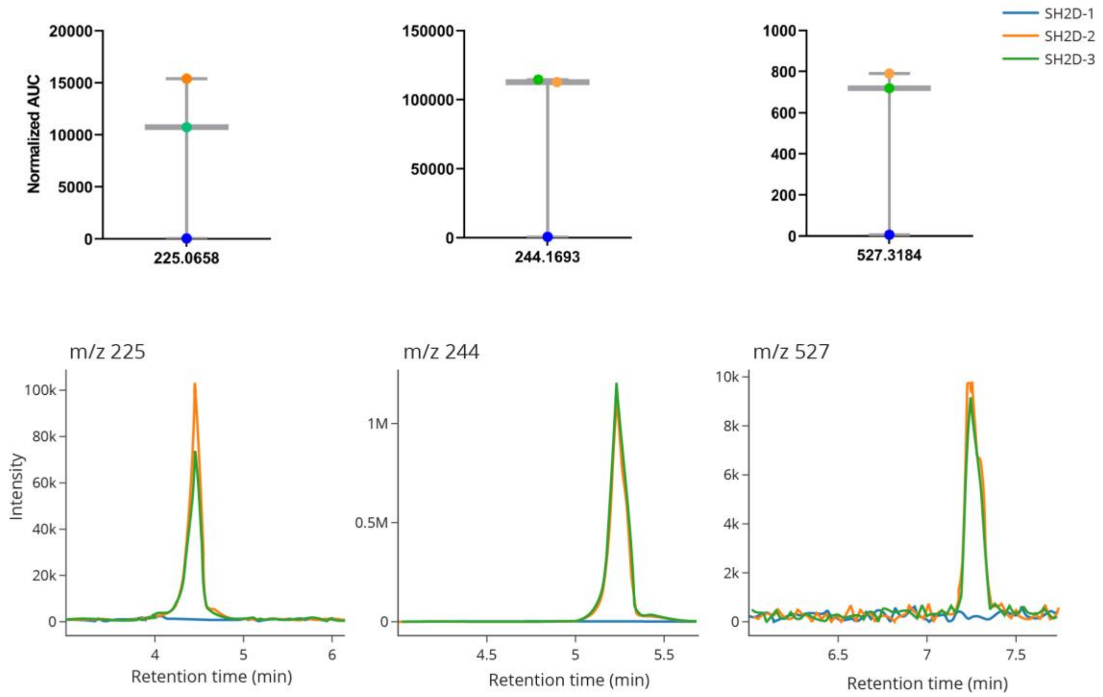


**Supplemental Figure 10. Using GNPS Dashboard to confirm that low abundance data is due to poor extraction or improper injection.** TOP: Boxplots with data points indicating the area under the curve value of features corresponding to *m/z* 225.0658, 244.1693, and 527.3184 of three biological replicates of *P. aeruginosa* SH2D overlaid on the boxes. BOTTOM: Extracted ion chromatograms of *m/z* 225.0658, 244.1693, and 527.3184 corresponding to each replicate.

Visualization of quantitative LC-MS data using boxplots or scatterplots can reveal outlier values. Due to the number of processing steps involved to generate quantitative values from raw LC-MS data, those outlier values may originate from poor sample extraction, improper injection volume, or inappropriate feature finding settings for the dataset. However, the GNPS Dashboard can be used to quickly visualize the raw data to identify the origin of the outlier values. For the example in **Supplemental Figure 10**, we analyzed the metabolome of *Pseudomonas aeruginosa* strain SH2D grown in synthetic cystic fibrosis medium 2 (SCFM2) containing dialyzed bovine submaxillary mucin (BSM). We subjected the LC-MS data to feature finding and generated box plots to visualize the area under the curve values for each replicate (**Supplemental Figure 10 TOP**). For many of the features quantified, biological replicate 1 (SH2D-1) values were near zero, while values for replicate 2 (SH2D-2) and 3 (SH2D-3) were not. This result suggested that there was likely an issue with either sample quality or data processing. We then visualized the extracted ion chromatograms (EIC) of *m/z* values associated with select features (**Supplemental Figure 8 BOTTOM**). Overlaying the EIC from each replicate for each *m/z* revealed that the boxplots faithfully represented the raw data, indicating an experimental issue associated with either sample extraction or injection.

GNPS Dashboard Link

**Use case 9: Cross Laboratory/Instrument sharing/visualization of MS data**

**Part 1: Multiple instruments within different laboratories**

**Monica Thukral - PhD Student - Microbial Ecology**

*Pseudo-nitzschia australis (P. australis*) is a cosmopolitan marine microalga capable of producing a potent neurotoxin domoic acid, particularly when undergoing nutrient starvation[23]. Domoic acid (DA) bioaccumulates up the food chain and can cause mass mortality to marine mammals and birds and also to humans if they consume DA-containing shellfish or crustaceans[23,24]. In the context of studying *P. australis'* metabolome under nutrient starvation, three LC-MS platforms are typically used by our group: A Bruker Amazon Ion Trap, an Agilent 6530 qToF, and a Thermo Fisher Scientific Q-Exactive qOrbitrap. As all three platforms come with their own vendor-specific data analysis software, the GNPS Dashboard facilitates this multi-laboratory and multi-instrument data analysis.

Visualization of mass spectrometry data of aqueous methanol extracts of *Pseudo-nitzschia australis are shown* in **SI Figure 11**. Panel a shows extracted ion chromatograms (XICs) (left) and mass spectra (right) extracted for m/z = 314.1441, the mass of domoic acid from ion trap (top) qToF (middle) and qOrbitrap (bottom). Due to the use of three different platforms for data acquisition, we expect the absolute value of the intensity of peaks to differ. To resolve this problem and ensure proper comparison, the XIC normalization toggle feature is turned on, to clearly and easily compare visual data. *A mass trace corresponding to domoic acids was clearly detected with all platforms and MS/MS spectra share the most intense fragment (m/z 266) but differ especially in the low m/z range in the ion trap data.* **SI Figure 11b** shows the comparison of data of extracts acquired on the Ion Trap and the two high-resolution qTOF and qOrbitrap platforms. Depicted are heat maps and total ion chromatograms (TICs) from qToF (left) and Orbitrap (right) provide a direct comparison of globe data structure, including MS/MS events. The heat maps show *m/z* vs retention time as a 2D LC-MS map. Blue crosses represent the triggering of MS/MS data collection. While in the qToF data, the blue crosses are denser, the qOrbitrap data shows blue crosses well distributed across the data. Also clearly visible is that dynamic exclusion is not utilized in the qToF analysis, causing multiple MS/MS to be triggered along one peak and reducing the metabolome coverage of a molecular network in downstream analysis. Hence, it is not surprising that the coverage of chemical space in the downstream molecular network data analysis has a significantly higher contribution from the qOrbitrap data (**SI Figure 11c**) than from the qToF.

In summary, the GNPS Dashboard directly benefited the investigation into *P. australis* metabolomics by allowing for rapid and efficient comparison between three instruments with different data analysis platforms. The GNPS Dashboard will serve as a valuable data integration tool for analysis going forward.

Classic Molecular Networking Link GNPS Dashboard IonTrap GNPS Dashboard qToF GNPS Dashboard Orbitrap

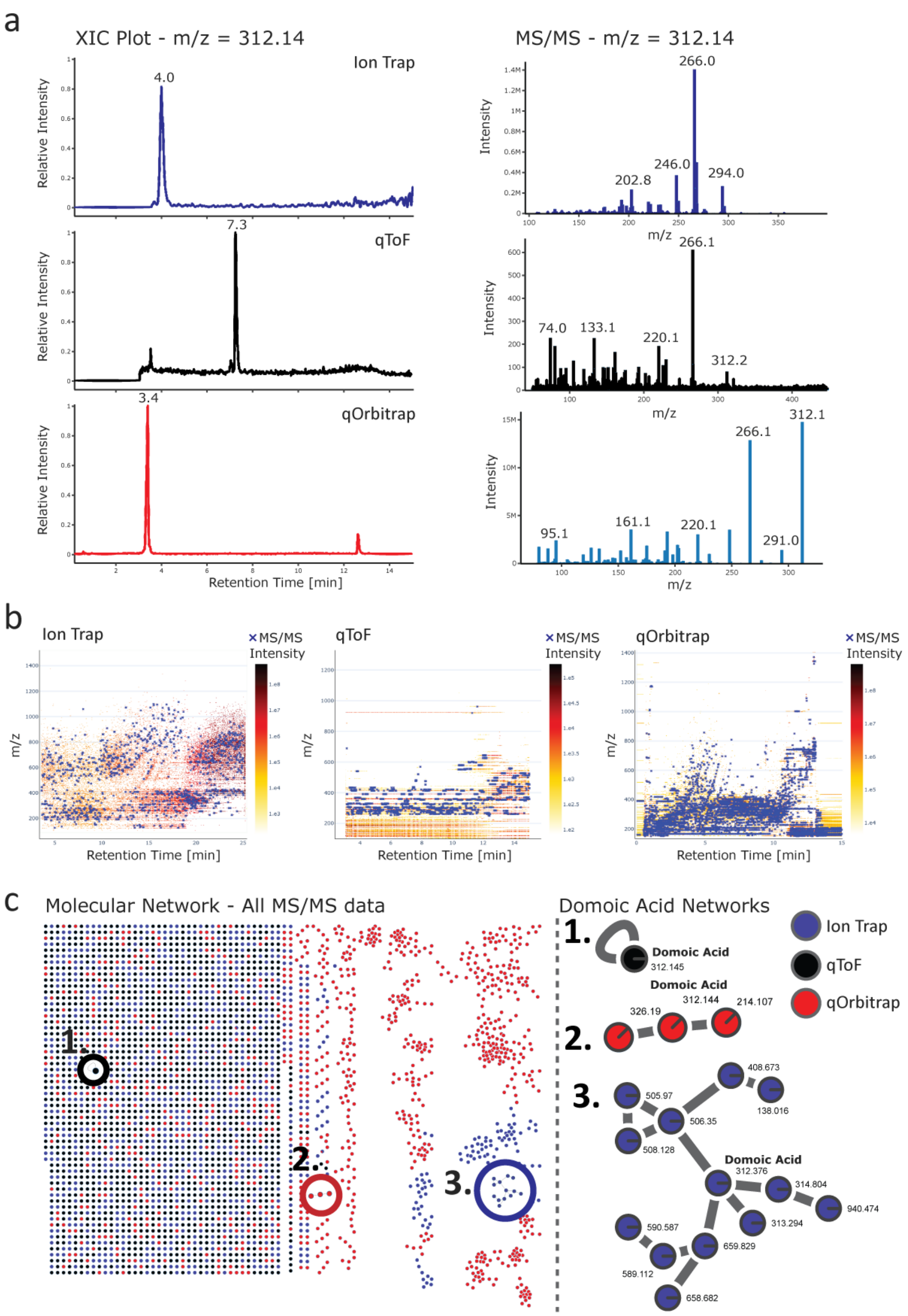


**Supplemental Figure 11: Cross-platform comparison of LC-MS/MS from the same sample**. Panel a shows the comparison of MS data from Domoic acid From extracts from *Pseudo-nitzschia australis* extracts that were recorded on three different instrument platforms (Ion Trap qToF and qOrbitrap). On the left side, the Extracted Ion Chromatograms (XIC) and corresponding MS/MS of domoic acid spectra on the right can be seen. Platform comparison of global DDA MS/MS data is shown in Panel b as heatmaps of the LC-MS run and MS/MS scans indicated as blue crosses. A molecular network from the global MS/MS data from all three platforms and zoom-ins to the domoic acids networks is shown in panel c.

**Part 2: Multiple instrument within the same laboratory**

**Alan Jarmusch PhD - Post-Doctoral Scholar - Analytical Chemistry**

The multitude of mass spectrometers in existence is reflective of their commonplace in modern chemical measurement as well as their flexibility. Different mass spectrometers are defined, principally, by the mass analyzer of which there are traps, time-of-flights, quadruples, sectors, and hybrids. The different analytical performances (*e.g.* sensitivity, specificity, speed, duty cycle, mass resolution) of these mass analyzers often necessitate multiple mass spectrometers in a laboratory in order to access the different types of experiments available. The practical implication of this fact is that a laboratory is often forced to purchase instruments from different manufacturers and learn each manufacturer's control and data analysis software. The data analysis software is often complex to learn, knowledge does not readily translate from one manufacturer's software to another, and is plagued by different file formats. In reverse order, the utilization of an open-source file format (e.g. .mzML[25]) is a good initial step toward overcoming this challenge. By learning a single data analysis and visualization software, particularly one that exists in a web browser, it is hugely advantageous and nearly eliminates the need to learn multiple pieces of manufacturer software for simple visualizations and data processing. This of course will not immediately eliminate the need for manufacturer software, but it does make the initial steps towards a democratized, open-source, web-enabled, trans-manufacturer, and trans-mass spectrometer piece of software to facilitate chemical measurement and minimize time spent learning disparate software.

**Use Case 10: Integration of Dashboard with other tools (SIRIUS) and comparison with standard to identify isomerization/degradation**

**Jessica M. Deutsch - PhD Student - Chemistry**
**Neha Garg PhD - Assistant Professor - Natural Product Chemistry**

In non-targeted metabolomics experiments, one can typically annotate 1- 10% of the detected metabolites. To enhance our ability to decode the dark metabolome, we utilize various *in silico* methods such as SIRIUS 4[26] with CSI:FingerID[27] among others. The use of GNPS Dashboard to generate XIC on chemical formulas suggested by SIRIUS followed by feature detection and interrogation of the presence of chemical formula across samples allows us to integrate output from tools such as SIRIUS 4 with CSI:FingerID with rapid inspection of $MS^1$ and $MS^2$ spectra across multiple samples. Analysis performed directly via the GNPS Dashboard eliminated the need to install and toggle between multiple software to extract features and visualize multiple XIC alongside SIRIUS 4 with CSI:FingerID. The option to generate XICs from molecular formulas in the GNPS Dashboard also enables automated visualization of potential adducts, such as $[M+Na]^+$ and $[M+K]^+$, which enables further validation of the proposed chemical formula. Furthermore, access to individual LC-MS files on GNPS Dashboard also enables visualization of the $MS^2$ spectra to inspect the fragmentation pattern and validation with analytical standards. Lastly, we can share these results with collaborators, additional lab members, and generate publication-quality figures without the use of expensive vendor software.

The use of GNPS Dashboard has been instrumental in a remote teaching setting where I teach a potpourri of multiple mass spectrometry-based data annotation strategies and has enabled remote training of graduate students that joined my laboratory during the COVID shut down. Thus, the advantages of GNPS Dashboard are multi-fold and extend beyond the research setting. In the example presented in **SI Figure 12**, we interrogated the annotation of korormicin A as suggested by SIRIUS with CSI:FingerID in organic extracts of a bacterium isolated from the coral tissue by our collaborator, Dr. Valerie Paul. The chemical formula was suggested as $C_{25}H_{39}NO_5$ by SIRIUS 4 with a score of 99.95% and the top-most annotation belonged to korormicin A (**SI Figure 12a**). The XIC for this chemical formula was generated in the organic extract of bacterium (**SI Figure 12b**), which revealed detection of both $[M+H]^+$ and $[M+Na]^+$ adducts. This feature was used to optimize parameters for MZmine-based[19] feature detection directly in GNPS Dashboard. These parameters were used to interrogate the presence of korormicin A in extracts of various microbes extracted from the same coral to identify likely isolation of the same species of bacteria (**SI Figure 12c**). Lastly, XIC comparison with analytical standards using GNPS Dashboard allowed us to observe isomerization of the purified molecule, which was also observed at low abundance in the culture extracts (**SI Figure 12c**).

GNPS Dashboard Figure 12b - Link
GNPS Dashboard Figure 12c - Link

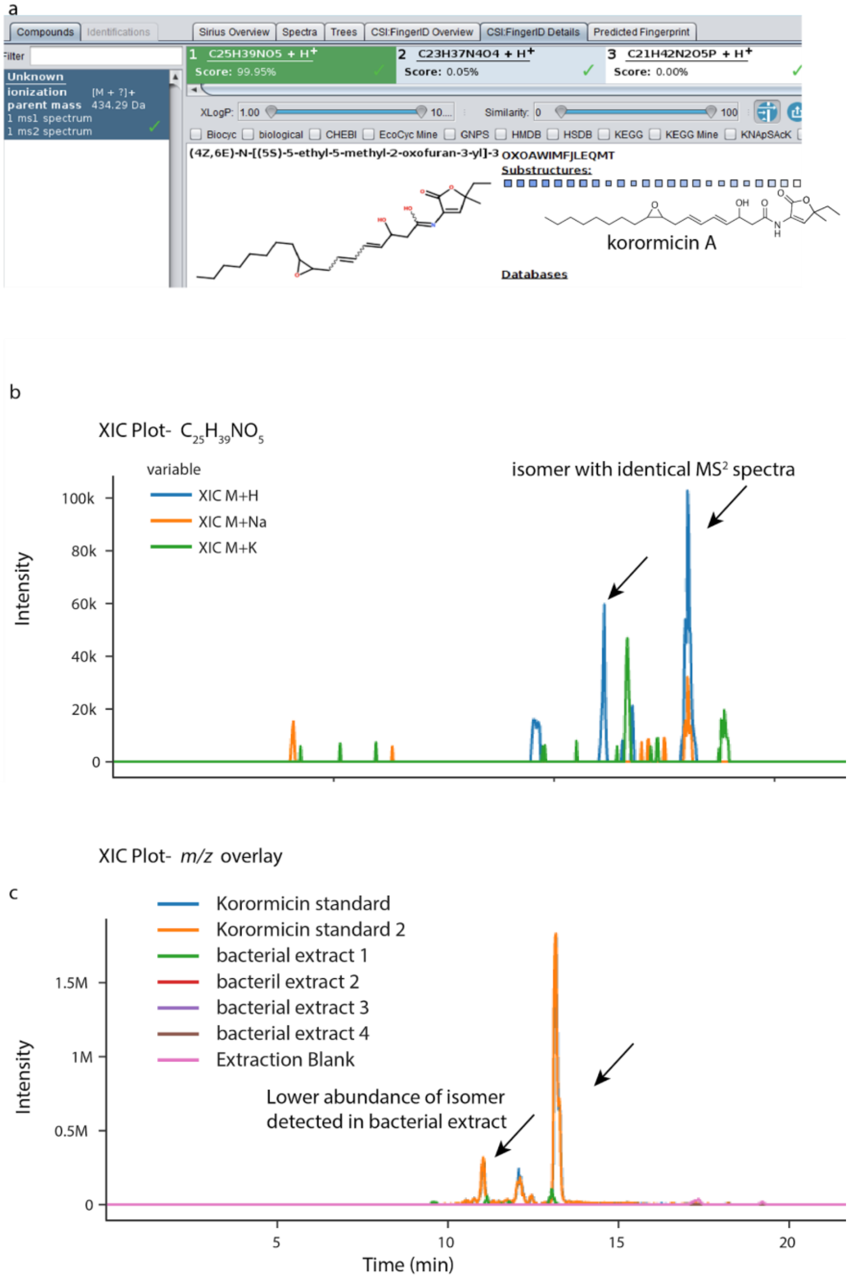


**Supplemental Figure 12. Integration of *in silico* annotation tool SIRIUS 4 with CSI:FingerID with GNPS Dashboard.** a) The output of SIRIUS 4 with CSI:FingerID showing prediction of chemical formula as $C_{25}H_{39}NO_5$, with plausible annotation as korormicin A as the top candidate for *m/z* 434.290 Da. b) The XIC plot for a chemical formula using LC-MS viewer shows the possible adducts as $[M+H]^+$ (blue trace) and $[M+Na]^+$ (orange trace). The trace for $[M+K]^+$ does not show peaks at the same retention time as $[M+H]^+$ confirming $[M+K]^+$ is not detected. c) The multiple XIC plot reveals that only one bacterial extract (green trace) produces korormicin A suggesting that the isolated bacteria is different from the other isolates. The same two isomers are observed in the bacterial extract. The isomer at 11 min is present in lower relative abundance in the bacterial extract than the standard suggesting conversion between the isomers during purification.

### Use Case 11: Targeted dereplication and evaluation of culture conditions

**Scott Jarmusch PhD - Post-Doctoral Scholar - Natural Product Chemistry**

Dereplication remains one of the principal target areas for metabolomics-based tools to accommodate the natural products community. GNPS molecular networking integrates fast, automated dereplication into its workflow, yet often, results still require the need to evaluate raw data for more critical interrogation. Utilizing the public data from a recent study on desferrioxamine metabolites produced by *Streptomyces* sp. S29[28], we can visualize how a targeted dereplication approach may be carried out. Using hits from the GNPS library as well as the work done by Traxler et al.[29], the GNPS Dashboard allows for fast evaluation regarding the presence or absence of accurate mass XIC as well as providing context in which samples these metabolites are present. Taking desferrioxamine D (*m/z* 603.37), we expect it to be produced and it is, in any of the three conditions tested (**SI Figure 13a**): monoculture, coculture with *Aspergillus niger* and coculture with *Botrytis cinerea*. Furthermore, in microbial natural products, evaluation of culture conditions is vital to gain a better understanding of metabolite elicitation, especially when focusing on discovery. The two studies above observe the production of long-chain acyl desferrioxamines exclusively under coculture conditions. When we evaluate the presence/absence of three acyl desferrioxamines ($C_{11}$-$C_{13}$) in the fungal coculture study, we can clearly see the presence of the three metabolites in the cocultivation with *B. cinerea*, a presence of one metabolite in cocultivation with *A. niger* and an absence of the three in monoculture conditions (**SI Figure 13b**). Beyond the evaluation of knowns, we can also easily visualize and allow for prioritization of extracts when hunting for unknowns. An unknown desferrioxamine-like metabolite was identified in this study and we observe that its presence is related to cocultivation with *B. cinerea*, providing a guide to future isolation of this unknown (**SI Figure 13c**). Allowing for this rapid observation in the GNPS Dashboard interface that can then link into further GNPS tools makes this functionality invaluable when analyzing data that would normally require multiple software types to be opened simultaneously, requiring high computing power that some researchers do not have access to. These basic functionalities shown are essential in the community and the ability of the GNPS Dashboard to facilitate these are paramount to the service it provides.

GNPS Dashboard Figure 13a - Link
GNPS Dashboard Figure 13b - Link
GNPS Dashboard Figure 13c - Link

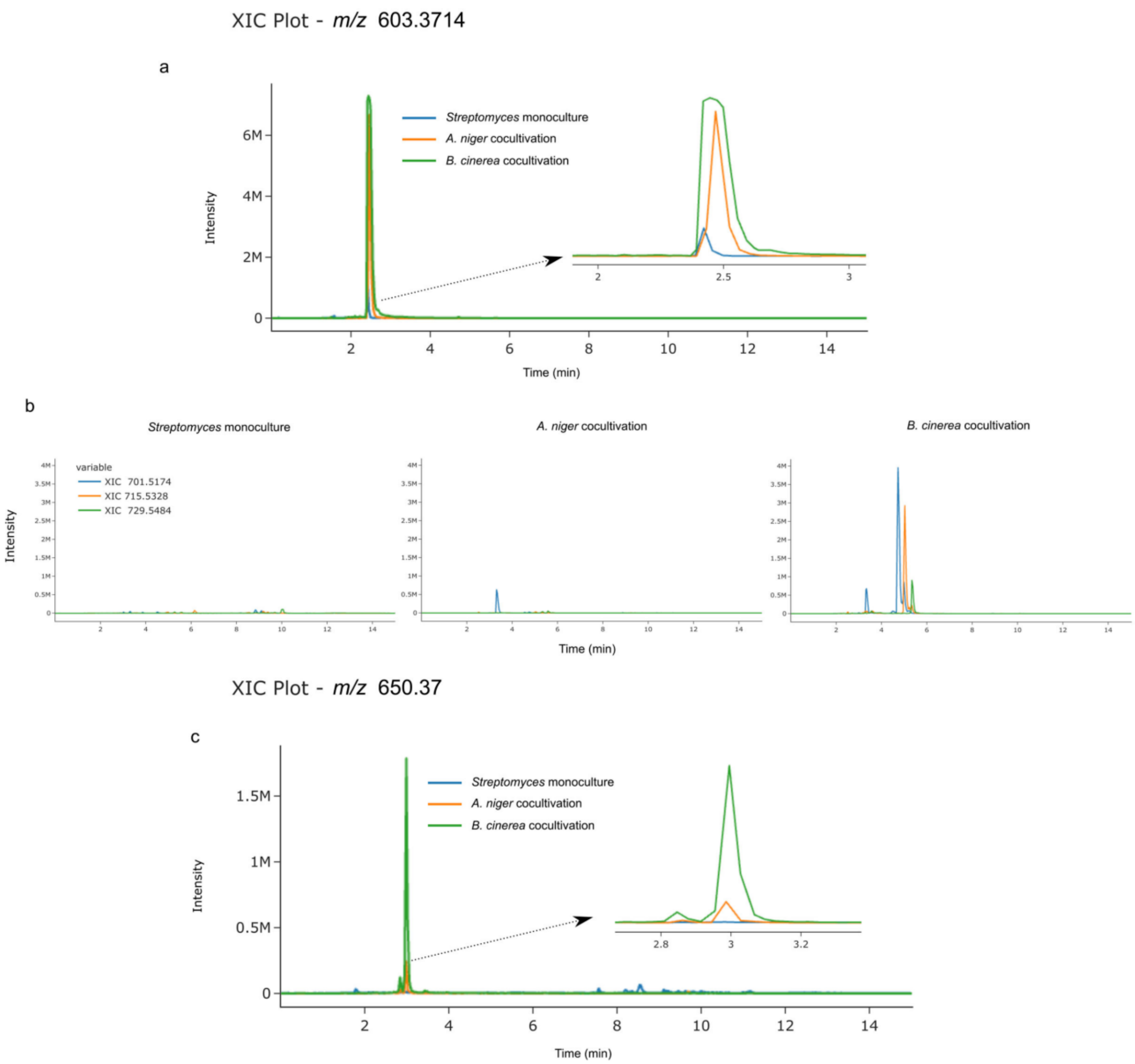


**Supplemental Figure 13. Targeted dereplication and evaluation of culture conditions.** a) XIC plot of desferrioxamine D (*m/z* 603.3714) using LC-MS viewer shows the presence of the archetypal siderophore in monoculture (blue trace), cocultivation with *A. niger* (orange trace), and cocultivation with *B. cinerea* (green trace). b) XIC plot grouped by file showing the presence/absence of acyl desferrioxamines C11-C13 in each culture environment, clearly showing production in the presence of *B. cinerea*, some production in the presence of *A. niger,* and no production in the monoculture. c) targeting the culture conditions where an unknown metabolite (*m/z* 650.37 $[M+2H]^{2+}$) could be isolated for future studies.

### Use Case 12 - Large Scale XIC Inspection in GNPS Dashboard

**Morgan Panitchpakdi - Staff Research Associate - Metabolomics**

Analysis of non-targeted LC-MS data often requires a thorough examination of the quality of each data file generated. Generating Extracted Ion Chromatograms (XIC) for internal standards and compounds of interest using vendor software is possible but becomes slower as the project size increases (>20 files). The GNPS Dashboard helps alleviate these issues and enables quick inspection and analysis of LC-MS data and can generate XICs for large-scale projects within minutes.

With the goal of reanalyzing published data from a clinical study investigating drug metabolism[30], the GNPS Dashboard was used to quickly analyze LC-MS data quality and to confirm the presence of five reported internal standards in 140 urine samples (MSV000082493) before continuing on with downstream processing with MZmine[19]. A single sample file was first inspected for the presence of the internal standards to determine retention time range. XIC's for all five internal standards, Omeprazole-d3 (*m/z* 349.1409), Caffeine-d3 (*m/z* 197.0992), Midazolam-d4 (*m/z* 330.1112), Dextromethorphan-d3 (*m/z* 275.2198), and Dextrophan-d3 (*m/z* 261.2035) were generated for all 140 urine samples. To obtain the XIC for all sample files, 140 USI's for these files were input into the GNPS USI field, *m/z* value and RT range were specified for each internal standard and visualized by grouping XIC's by *m/z* (**Figure 14a**). The GNPS Dashboard was used to visualize plotted AUC values for each of the five internal standards in all 140 urine samples (**Figure 14b**). In summary, the GNPS Dashboard generated XIC's for 140 sample files faster than the vendor-provided data analysis software. Visual and shareable AUC value plots allow for quick analysis of internal standard presence and data quality before downstream processing and analysis begins.

GNPS Dashboard Link - Omeprazole-d3
GNPS Dashboard Link - Caffeine-d3
GNPS Dashboard Link - Midazolam-d4
GNPS Dashboard Link - Dextromethorphan-d3
GNPS Dashboard Link - Dextrophan-d3

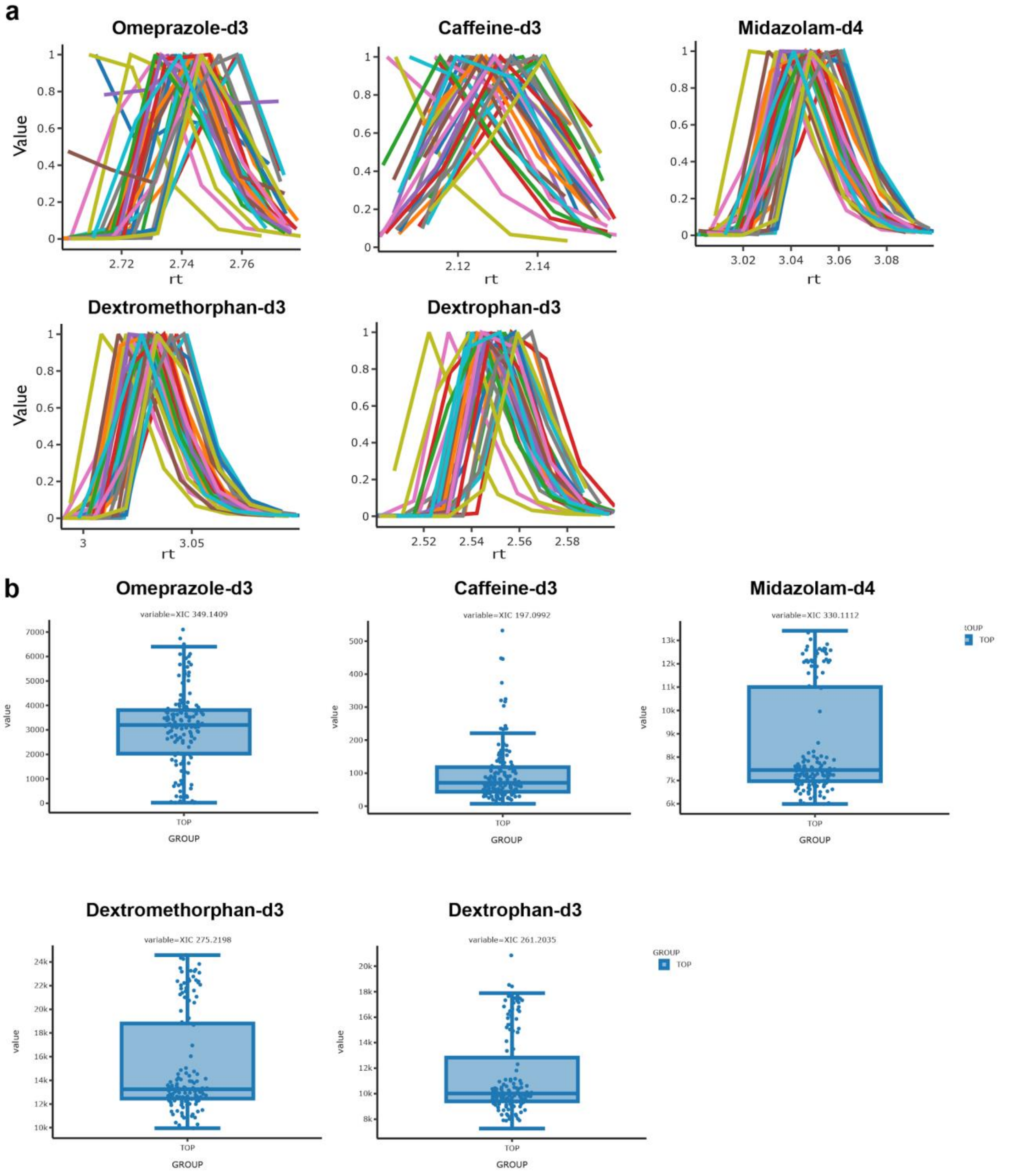


**Supplemental Figure 14. Large Scale XIC in GNPS Dashboard**

a) Ovelrayed Extracted Ion Chromatograms (XIC) for each of the five internal standards in all 140 urine samples b) Plotted AUC values to confirm the presence of five internal standards in all 140 urine samples.

### SI Use Case 13 - Validating Multiply-Eluting Peptides

**Ben Pullman - PhD Student - Computational Mass Spectrometry**
**Nuno Bandeira PhD - Professor - Computational Mass Spectrometry**

When analyzing a label-free proteomics experiment, the assumption is often made that each peptide should elute a single time in a given fraction, and the intensity of the peptide is that of the feature. However, in some cases, peptides can elute multiple times over a single run. Multiple elutions can complicate the calculation of feature intensity as well as create issues for the transfer of features between different runs. The GNPS Dashboard allows us to validate these multiply eluted peptides when examining the feature intensities.

In the following example (GNPS Dashboard link), from the recent experiment looking at 31 tissues of the human proteome[31] run through MaxQuant[17] for feature finding and identification, we see an example of a peptide:

NDGYLM[+16]FQQVPM[+16]VEIDGM[+16]K

mapping to protein group:

sp|P08263|GSTA1_HUMAN;sp|P09210|GSTA2_HUMAN

that elutes 10 times over a 30-minute span in a single LC-MS run. All of these features have a mapped MS2 that could be identified to the peptide as well. Further, we see that the intensity of all the features is at max only a single order of magnitude lower than the highest intensity, and if the features beyond the highest two are not considered, the total intensity for the peptide would be off by ~30%.

Using the GNPS Dashboard, we can examine the MS2 spectra for each feature in the XIC, and see that while there are likely some co-eluting peptides, the MS2 spectra generally look similar, confirming that indeed this feature likely elutes multiple times. The USIs for these MS2s are below:

mzspec:MSV000083508:01275_D03_P013127_S00_N20_R1:scan:24647:NDGYLM[+15.994915]FQQVPM[+15.994915]VEIDGM[+15.994915]K/2
mzspec:MSV000083508:01275_D03_P013127_S00_N20_R1:scan:30728:NDGYLM[+15.994915]FQQVPM[+15.994915]VEIDGM[+15.994915]K/2
mzspec:MSV000083508:01275_D03_P013127_S00_N20_R1:scan:35674:NDGYLM[+15.994915]FQQVPM[+15.994915]VEIDGM[+15.994915]K/2
mzspec:MSV000083508:01275_D03_P013127_S00_N20_R1:scan:41691:NDGYLM[+15.994915]FQQVPM[+15.994915]VEIDGM[+15.994915]K/2

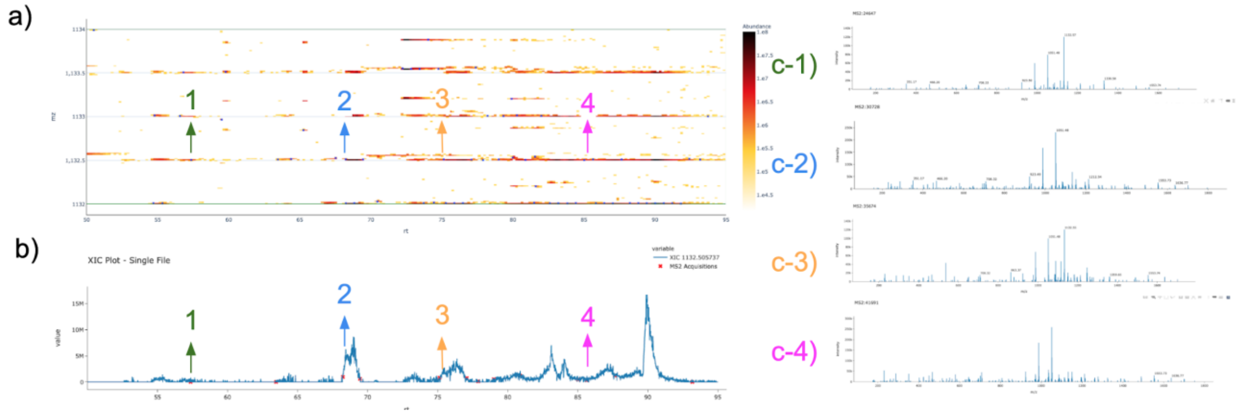


**Supplemental Figure 15**: **Example of a long eluting peptide**

a) Shows the LC-MS map for the features of NDGYLM[+16]FQQVPM[+16]VEIDGM[+16]K zoomed in to show only RT 50-95 min and *m/z* 1132-1134. b) Shows the XIC for the mono-isotopic signal of the feature (*m/z* 1132.505737) with a retention time of 50-95 min, c) 1-4 show the corresponding MS/MS spectra that map to the identified features, color-coordinated with the arrows showing where the MS/MS was acquired on both a) the LC-MS map and b) the XIC.

### SI Use Case 14 - Implementation of third party tool in the GNPS Dashboard for rapid prototyping

**Robin Schmid PhD - Post-Doctoral Scholar - Computational Mass Spectrometry**

Software solutions for mass spectrometry data analysis have become essential to cope with the complex data generated by non-targeted LC-MS/MS. The GNPS dashboard offers collaborative visualization and integrative data analysis options (e.g., feature finding, molecular networking, and repository scale spectrum searches). Beyond a visualization tool for end-users, the GNPS Dashboard provides a new platform for software developers to quickly test new MS data analysis tools and make them readily accessible to the broader community. The access of public metabolomics and proteomics data in mass spectrometry repositories, rich visualization options, and the GNPS Dashboard's web-interface are beneficial for the testing and development of new software tools. Open-source licensing and effortless local deployment of the GNPS Dashboard enable rapid prototyping to integrate other tools and tailor the modular dashboard with specific graphical and tabular outputs. To exemplify a use case for developers, we used the GNPS Dashboard to integrate and test the latest development branch of the feature finding tool MZmine 3, which is currently undergoing a comprehensive redesign of its data model. The goal was to provide a framework to deploy, run, and visually compare feature finding results from MZmine 3 and the latest release of MZmine 2 as a consistency check.

After downloading the open-source code for GNPS Dashboard, the dashboard can be executed locally in debugging mode (Documentation). MZmine 3 was integrated based on the existing feature finding options. Instead of running MZmine 2 only, both versions were executed sequentially to yield feature tables. Here, the dashboard provided UI controls to set essential parameters (e.g., mass tolerance, noise level, and feature width constraints) and rerun the comparison. New parameters that were introduced in MZmine 3 were set to reflect the default behavior of MZmine 2 (e.g., the minimum number of data points in the local minimum search feature resolver). We adjusted the LC-MS 2D heatmap (**Supplementary Figure 16**) and feature finding results table to overlay the results of both MZmine versions.

The presented integration was beneficial for comparing the latest development version and release of MZmine on different metabolomics datasets. As a result, we were able to track behavior changes in data processing and reporting. Some identified as planned changes (e.g., new parameters), while others resulted from introducing a new data model (e.g., changed accuracy (double<>float) for various data types) or changes in calculation methods. The latter case led to the feature area being calculated based on the retention time in minutes instead of seconds. The dashboard helped identify these small differences and verify that feature finding resulted in the same number of features for both MZmine versions. A discrepancy in the number of detected features was evident based on missing shapes in the LC-MS map for MZmine 3 when a new filter parameter was not turned off (**Supplementary Figure 16**).

Even for a rich graphical user interface tool, such as MZmine, the integration into the LC-MS dashboard facilitated the software development and testing. Key points were the provided data access from various repositories, the LC-MS map that showed overlapping or missing features, and the integration of MZmine 3 into the dashboard's modular framework. Furthermore, the dashboard's deployment in a local development environment was effortless and followed a few documented steps (Documentation) only.

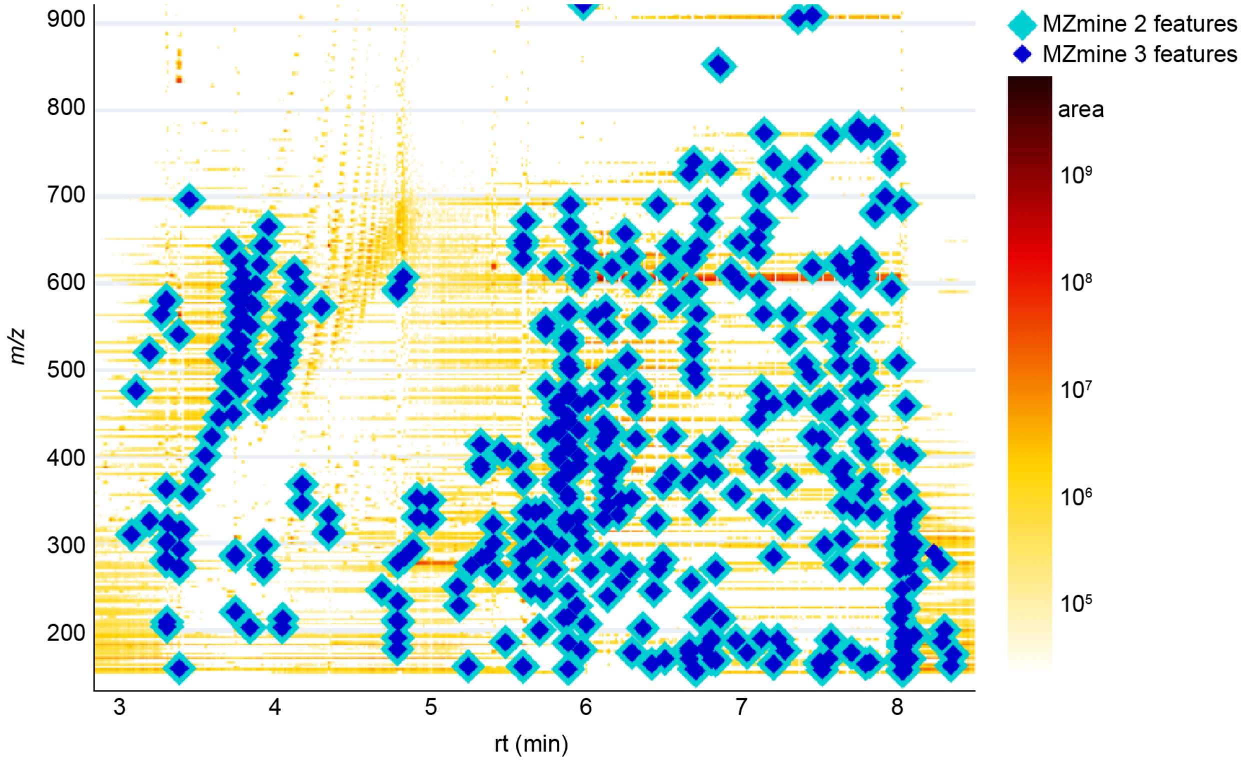


**Supplemental Figure 16: Consistency testing by overlaying feature finding results from MZmine 2 and the latest development version of MZmine 3.** A complete results overlap was achieved after adjustment of the MZmine 3 preferences and batch mode. The specialized version of the GNPS dashboard informed the development of MZmine 3 to provide comparable results to MZmine 2.

### SI Use Case 15 - Quantification of isobutanol produced by metabolically engineered *E.coli* strains

**Simon Boecker, PhD - Post-doctoral Scholar - Metabolic Engineering**

The performance of engineered microbial cell factories for the synthesis of a product of interest is usually characterized by the three parameters titer (product per volume), productivity (product per volume (or per biomass) and time), and yield (product per substrate). To determine these parameters, an exact quantification of the produced metabolite secreted to the medium is indispensable.

For this use case, we compared the performance of two *E. coli* strains, which were engineered for the production of the biofuel isobutanol (*i*BuOH). *i*BuOH is a volatile compound and can be quantified directly from the culture broth by GC-MS using head-space injection. External standards and medium sampled at different time points were analyzed on an Agilent Technologies GC-MS instrument in SIM mode. In our lab, we only own one license of the MassHunter software for data analysis, complicating the evaluation of the obtained data on your office or home PC. Instead of using MassHunter, we converted the data to mzML-format via the MSConvert tool[14], uploaded it to the MassIVE repository, and used the GNPS Dashboard for data analysis. XICs were extracted for the quantifier ion (*m/z* 56.1) and the areas of the selected peaks were automatically integrated and exported as csv-file and used for generating a calibration curve (**SI Figure 17a**). The amount of *i*BuOH produced by the two strains was quantified with the calibration curve, demonstrating that strain 2 shows slightly superior performance in terms of *i*BuOH production (**SI Figure 17b**).

In summary, the GNPS Dashboard is a very useful tool that facilitates data analysis for compound quantification by GC-MS, especially when the number of available software licenses is limited in the lab.

GNPS Dashboard integration results - Link
GNPS Dashboard Figure 17 - Link

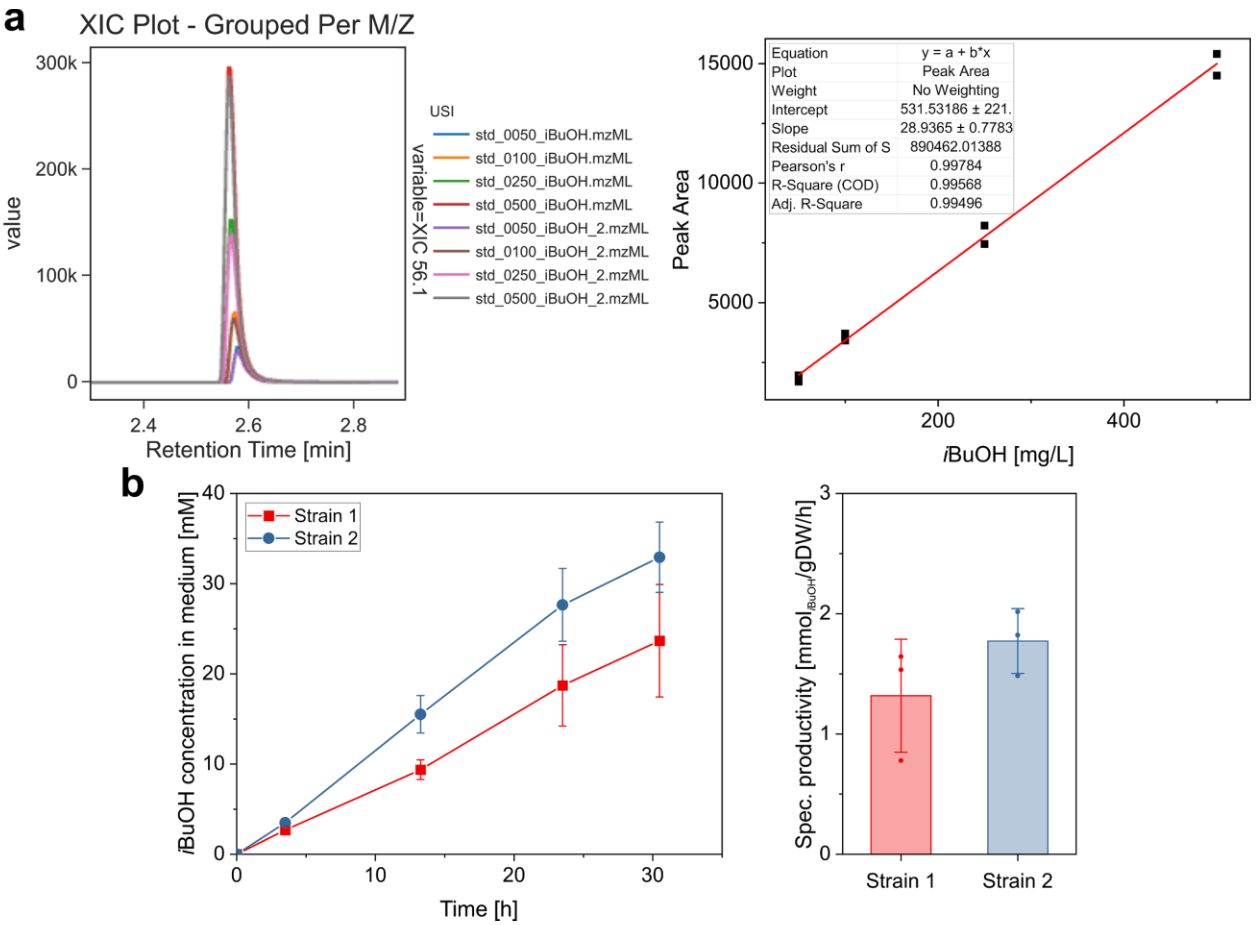


**Supplemental Figure 17:** a) Overlaid XICs for *m/z* 56.1 of *i*BuOH standards (in duplicates) visualized in the GNPS Dashboard and generated calibration curve from integrated peak areas. b) Time course of *i*BuOH production and specific productivity of both *E. coli* strains.

### SI Use Case 16 - Exploring new directions in previously-analyzed GC-MS data

**Rachel Gregor, PhD - Post-doctoral Scholar - Chemical Ecology**
**Michael M. Meijler, PhD - Professor - Chemical Biology**
**Itzhak Mizrahi, PhD - Professor - Microbial Ecology**

Large-scale ‘omics projects generate vast quantities of mass spectrometry data, more than can be comprehensively analyzed within the scope of one or even a series of publications. The same data can often be re-analyzed later to explore different questions or angles, especially if it is uploaded on public data depositories such as MassIVE. For example, as part of a large fecal microbiome-metabolome dataset collected in collaboration with a local zoo, we included food samples from the animals’ diets and analyzed them by GC-MS. The food GC-MS metabolomes were not characterized in detail in the original data analysis, other than to quantify the overlap between fecal and food metabolites. Therefore, this portion of the dataset would be an excellent candidate for future analyses, for example in order to explore specific classes of nutrients found in different animal diet components.

However, unexpected setbacks can arise when re-analyzing data at a later point, even for the original team who collected the data. In this case, we had originally visualized the chromatograms using Agilent’s proprietary MassHunter software and performed deconvolution and molecular networking using the GNPS MS-Hub workflow. Since then, the lead author in this project finished her PhD and moved to a different institution, and no longer has access to an Agilent workstation with MassHunter, presenting a barrier in visualizing the chromatography data. This stage is especially crucial for the analysis of GC-MS data with electron impact ionization, both in order to manually verify the quality of the deconvolution for peaks of interest, and because different classes of compounds often have characteristic retention time ranges that are evident on the chromatograms.

Using the GNPS Dashboard, we were able to load the GC-MS dataset from MassIVE (MSV000083859) and immediately explore metabolites of interest in the food samples. For example, the herbivores in the study consumed a variety of foods, including fresh fruits and two types of pellets. We examined the sugar content of three types of fruit and the two pellet samples by creating an XIC for m/z 217, a characteristic sugar fragment, resulting in three groups of peaks corresponding to mono-, di-, and tri-saccharides (**SI figure 18a**). It was immediately evident that fruits were enriched in mono- and some di-saccharides, while the pellets contained primarily di- and tri-saccharides. This was further strengthened by the molecular network, which showed a similar trend in a large cluster of sugar compounds (**SI figure 18b**). Using the GNPS Dashboard and the identifications from the molecular networking, we were able to easily examine specific examples of sugars from each type, such as the mono-saccharide fructose, the di-saccharide sucrose, and the tri-saccharide maltotriose, and compare the relative quantification in each sample (**SI figure 18c**). This analysis was possible quickly and easily upon selecting these samples from the MassIVE dataset on the GNPS Dashboard, thus removing barriers and making this data immediately accessible and ready for analysis.

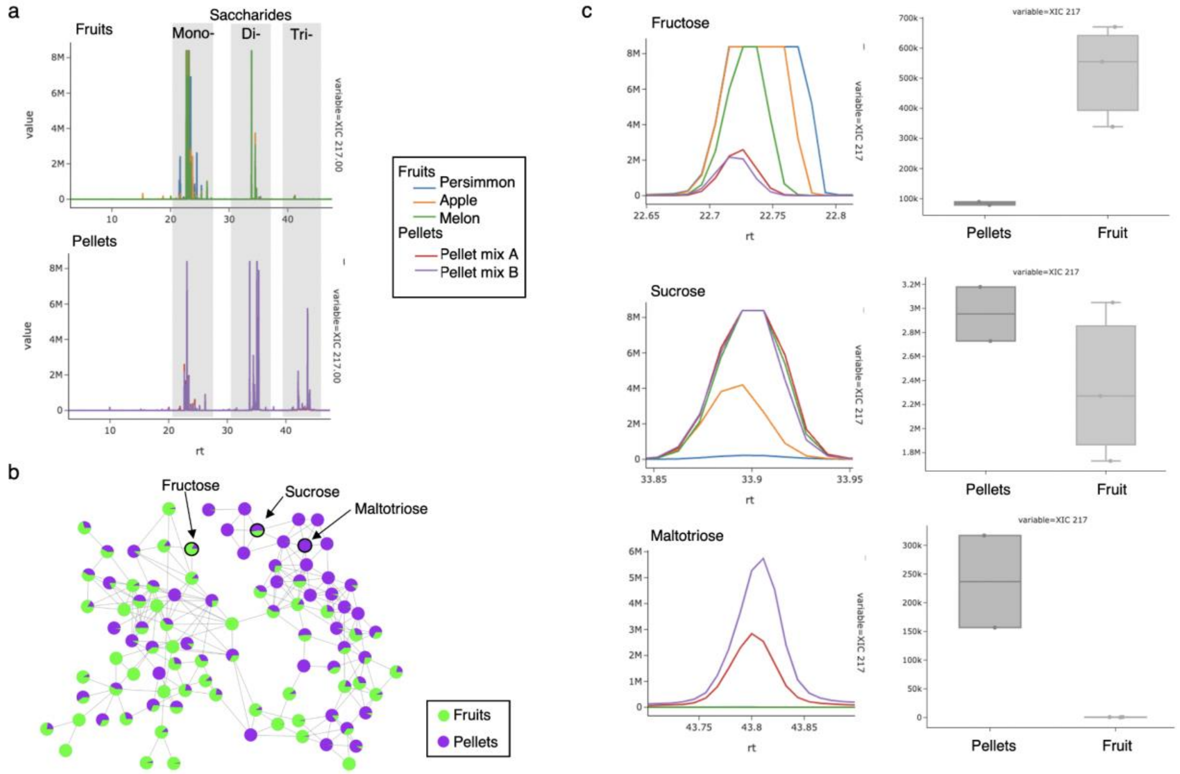


**Supplemental figure 18.** a) Overlaid XIC's for m/z 217 for fruit samples (top) and pellet samples (bottom). Characteristic elution ranges for mono-, di-, and tri-saccharides are indicated in grey. b) Molecular network of saccharide cluster with relative abundances of fruit (green) and pellets (purple). (c) Overlaid XIC's for m/z 217 for all samples, for peaks identified by spectral library search as fructose (top), sucrose (middle), and maltotriose (bottom).

### SI Use Case 17 - Inspection of data during the review process.

**Pieter C. Dorrestein, PhD - Professor - Small Molecule Mass Spectrometry**

The review process is an active process of the scientific enterprise. The goal of peer review of manuscripts is to ensure that the science that is presented is accurate and, when needed, to provide suggestions to improve the work. One of the key challenges as a reviewer of research articles that include mass spectrometry (MS) data is to get insight into the methods and settings used for data acquisition and quality or details of the actual data that is collected. Even when data is made public, many reviewers do not have or take the time to look at the raw MS data as there is a high barrier to entry (e.g., as this entails downloading and converting the data into the proper format and then loading the data into specific software to view). The GNPS Dashboard makes this process a simple single step in the web browser. Further,  if the authors include a direct url, then it is as easy as clicking the link to allow direct data inspection. As the review process is confidential, in this use case I will not show data, but rather describe recent examples that have been observed during recent reviews.

Being able to inspect the data during a review enables one to gain insight if there are alternative explanations possible. Most methods in papers have insufficient detail to understand how the data was acquired. A quick inspection of the *m/z* and retention time (rt) data with MS/MS points marked will reveal specific details about the data dependent acquisition (DDA), starting with the length of separation, how well the TIC appears to separate, general noise levels, how much data was also present in the background or blank samples, etc. Quick inspection using the GNPS Dashboard also gives insight into the use of data dependent exclusion parameters. For example continuous “lines" of MS/MS at a given m/z value, can reveal that dynamic exclusion was not employed or not properly set, and the DDA perhaps cycled through the top n most intense ions only. Being able to quickly inspect the data in this way will enable reviewers to rapidly judge if the methods and data are appropriate for the task presented in the paper. For example if the paper wants to get molecular networks from as many different molecules as possible then it is critical to set the DDA including dynamic exclusion appropriately, however if the task is to verify the presence of a few molecules and their analogs, other settings may be appropriate.

In addition, with the GNPS Dashboard  it is possible to quickly check how the data was used and if the data analysis is reproducible. For example, a paper may show an zoomed XIC derived feature but upon inspection by zooming out there are multiple XIC’s with the same mass. Thus, integration of peak area needs to be carefully considered as there could be artefacts due to a different molecule with the same mass (which will also give rise to a different MS/MS spectrum), an isomer which would likely have a nearly identical MS/MS spectrum, but could also be the same molecule but with different ion adducts, rotamers of molecules. In other words a single molecule can give rise to multiple peaks during chromatography. This could affect the quantification results and thus is important to be able to assess this as a reviewer. Being able to inspect the data will not only allow reviewers to inspect the details of the data and data acquisition parameters but also share their observations as hyperlinks in the reviews so that reviewers can help the authors to improve their papers or to provide suggestions on what to consider in future experiments. This way we all grow scientifically due to the review process.

**Use Case 18: Structure and fragmentation pattern analysis through the full collaborative synchronization mode**

**Mirtha Navarro PhD – Professor – Natural Product Chemistry, Bioactivity, and Nanotechnology**

**Felipe Vasquez-Castro – MSc candidate – Natural Product Chemistry and Environmental Microbiology**

Depending on the structure of research groups, principal investigators (PI) are directly involved in the day-to-day knowledge transfer to junior researchers. Therefore, tools for efficient collaborative work are of the utmost importance to support these interactions. For instance, the analysis of complex mass spectrometry data requires particular supervision, and this time-consuming effort becomes a challenge for Principal Investigators' tight schedules. GNPS Dashboard facilitates this task through the fully collaborative synchronization mode, which allows a multi-user simultaneous visualization and examination of data, thus enhancing the learning experience. Hereby we illustrate such a case, in which a PI and a junior researcher use GNPS Dashboard to perform a remote multi-user analysis of a LC-MS/MS spectrum with the purpose of identifying the structural differences and fragmentation patterns of procyanidin oligomers. The researcher needed to learn and apply these interpretation skills in proanthocyanidin-enriched extracts from the PI's research group (BIODESS). For this, the spectrum (MSV000087075) of a *P. domestica* enriched-polyphenolic extract obtained using Pressure Liquid Extraction and Amberlite resin chromatographic purification[32], was loaded onto the GNPS Dashboard, and the full collaborative synchronization mode under Sync Options was initiated (**SI Figure 19**).

During inspection of the Data Exploration panel, the PI highlighted that a peak at m/z 577.13 (12.64 min) indicated the presence of a procyanidin B-type dimer holding one single interflavanic bond between two units of (epi)catechin, while a peak at m/z 575.12 (24.96 min) indicated a procyanidin with 2 Da difference. Thus, corresponding to an A-type dimer with two single bonds present between the flavan-3-ol units. Based on this observation, the researcher was able to assign a peak at m/z 865.20 (17.88 min) to a procyanidin B-type trimer and to observe the peak at m/z 863.18 (18.80 min) with 2 Da lower mass. Therefore indicating a mixed procyanidin type-A trimer with one and two single interflavanic bonds respectively between the three (epi)catechin units. (**SI Figure 20**).

The utilization of the full collaborative synchronization mode focused afterwards in the PI explanation of the different fragmentation pathways these compounds undergo, namely Retro Diels-Alder (RDA), Heterocyclic Ring Fission (HRF) with the loss of phluoroglucinol (126 Da), and Quinone-methide (QM). The spectrum analysis using the GNPS Dashboard allowed the researcher to observe the more abundant ion fragment at m/z 425 in the procyanidin B-type dimer corresponding to the RDA neutral loss of 152 Da. The loss of water (18 Da) after RDA fragmentation produced the more abundant ion fragment at m/z 695 in the procyanidin B-type trimer. In addition, the joint spectrum analysis allowed the PI and researcher to highlight differences in respect to the second preferred fragmentation pathway. In fact, the second more abundant ion fragment at m/z 451 in the B-type dimer corresponded to the HRF pathway after the neutral loss of phloroglucinol (126 Da), while the QM pathway delivered the second more important ion fragment at m/z 577 in the case of the B-trimer (**SI Figure 21**).

In summary, GNPS Dashboard permitted the simultaneous exploration and analysis of mass spectrometry data through the fully synchronized collaborative mode, which proved useful for a Principal Investigator to share information in an efficient way with a researcher in MS data analysis, enhancing the learning process through the possibility of reviewing all actions and provide simultaneous feedback on the data produced.

Link to XIC: [GNPS - LCMS Browser (ucsd.edu)]
Link 2D LC-MS [GNPS - LCMS Browser (ucsd.edu)]
MS2 at 12.64 min: [GNPS - LCMS Browser (ucsd.edu)]
MS2 at 17.88 min: [GNPS - LCMS Browser (ucsd.edu)]

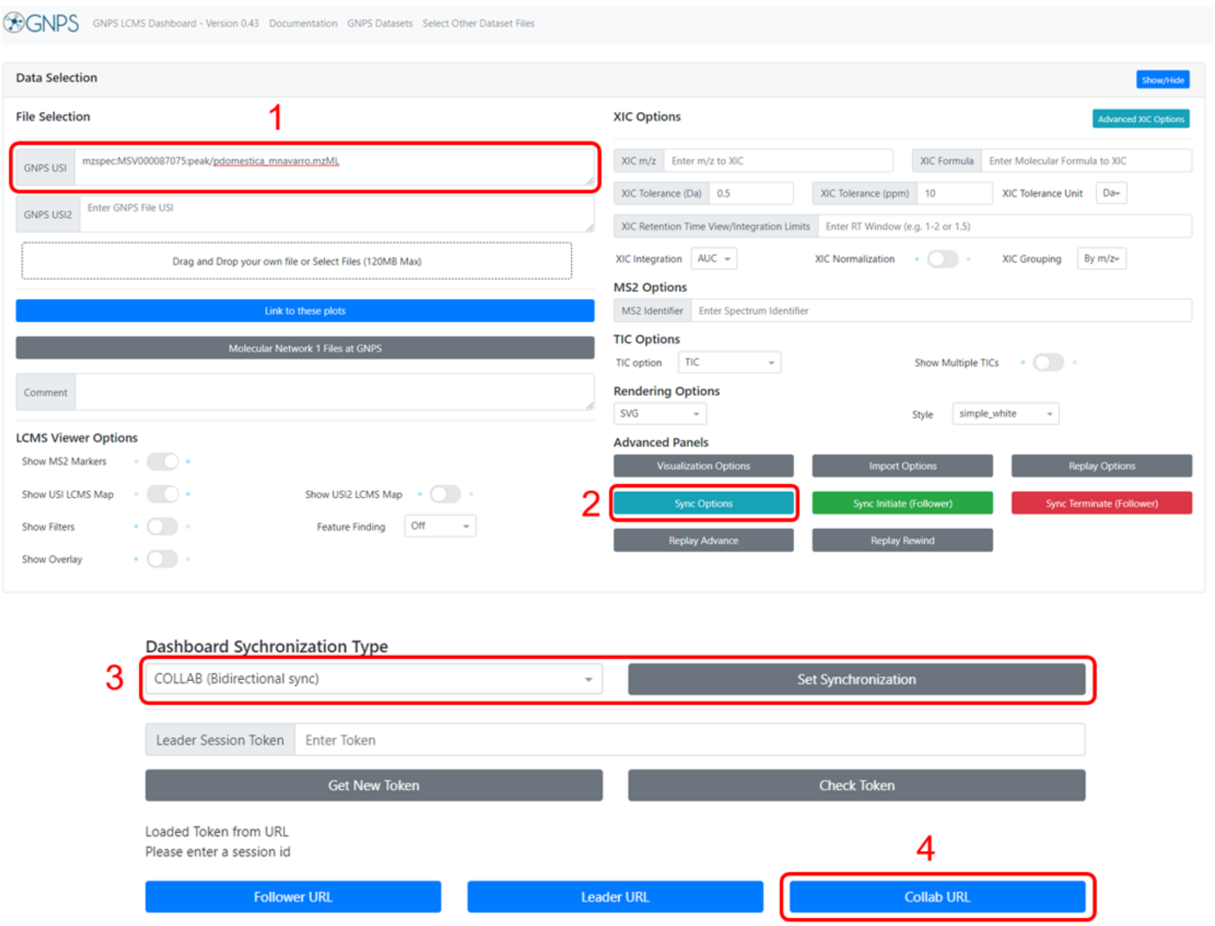


**Supplemental Figure 19**: Step-by-step setup of the bidirectional synchronization mode. A spectrum is loaded onto GNPS Dashboard using a MassIVE accession number (as in this case) or by dragging and dropping the file. The spectrum can either be loaded before or after initiating the bidirectional synchronization mode. The button ‘’Sync Options’’ reveals the different synchronization types available. The synchronization is set to ‘’Bidirectional sync’’ under ‘’Dashboard Synchronization Type’’, and the URL to the session is obtained by clicking on the ‘’Collab URL’’ button.

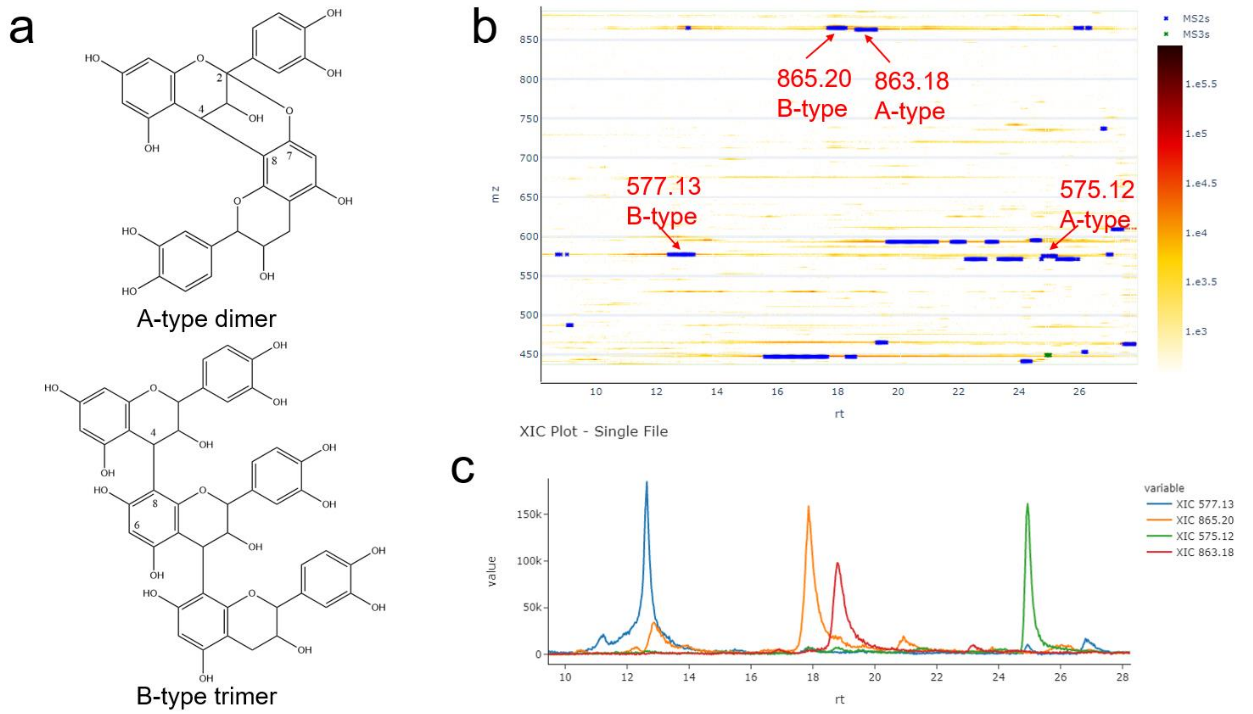


**Supplemental Figure 20**. a) Example structures of a procyanidin A-type dimer constituted by two units of (epi)catechin with C4-C8 and C2-C7 bonds and a procyanidin B-type trimer composed of three units of (epi)catechin with C4-C8 bonds. b) 2D LC-MS heatmap for procyanidin A-type and B-type dimers and trimers c) grouped XICs for procyanidin A-type and B-type dimers and trimers.

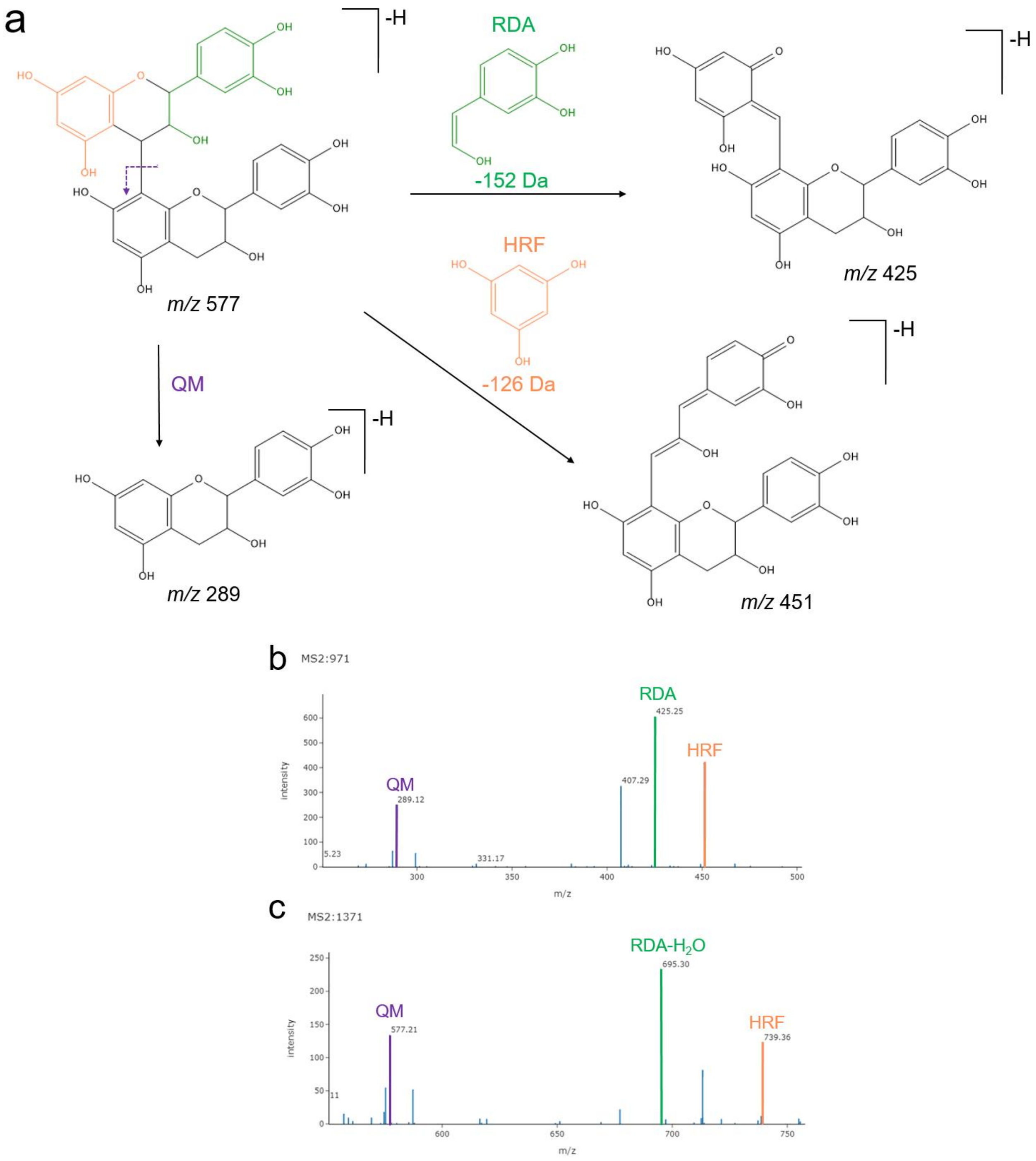


**Supplemental Figure 21**. a) Fragmentation pathways for procyanidins: Retro Diels-Alder (RDA), Heterocyclic Ring Fission (HRF), and Quinone-methide (QM). b) MS2 spectrum of a procyanidin B-type dimer at 12.64 min (m/z 577.13). c) MS2 spectrum of a procyanidin B-type trimer at 17.88 min (m/z 865.20). The fragments highlighted in both spectrums represent the respective fragmentation pathways illustrated in panel a.
